# Top-down semantic predictions align phonetic neuronal dynamics in human superior temporal gyrus

**DOI:** 10.1101/2023.10.30.564638

**Authors:** Pavo Orepić, Wilson Truccolo, Eric Halgren, Sydney S. Cash, Anne-Lise Giraud, Timothée Proix

## Abstract

Speech processing involves a hierarchy of cognitive layers from low-level phonetic to high-level semantic representations. However, interactions are not only feed-forward; Feedback mechanisms are crucial for real-time, accurate speech recognition. Yet how distant levels interface during speech processing remains unclear. Here, we analyzed intracortical recordings from 624 neurons across three human participants implanted with microelectrode arrays in the anterior superior temporal gyrus during an auditory semantic categorization task and natural speech perception. We identified distinct neural subspaces, or manifolds, for semantic and phonetic features, with a functional separation of the corresponding low-dimensional dynamics. We contrasted a bottom-up cumulative and a top-down predictive hypothesis on phonetic-semantic temporal alignment, and found phonetic alignment to word-level semantic representations, signaling top-down prediction. These effects were consistent across participants at the spiking level, and remained undetected in adjacent ECoG recordings. These findings demonstrate the reorganization of neuronal population dynamics supporting phonetic representations during semantic prediction.

## Introduction

Cognitive models of speech perception propose that speech sounds are processed into meanings by a series of neural modules. In the brain, semantic and conceptual representations emerge in the ventral stream (i.e. the ”what” stream) [1], where perceived speech is sequentially transformed from spectro-temporal encoding of sounds in Heschl’s gyrus, over phonetic features in the superior temporal gyrus (STG) and sulcus, to lexical and combinatorial semantics in the anterior temporal lobe (ATL) [2, 3]. The ATL is considered a specialized semantic module, connected to modality-specific sources of information [4, 5], with causal semantic impairments occurring after bilateral ATL damage [6–9]. Higher levels of semantic representations appear largely distributed at low temporal resolution [10], yet critically rely on the earlier steps of lexicosemantic processing. This modular and sequential perspective implies that a complex neural process transforms speech features from one functional brain region to the next up to lexical and multimodal conceptual representations.

Recent neuroimaging findings and AI models suggest, however, that these early transformations might be less modular and sequential than predicted by the classical hierarchical view of language processing [5, 11, 12]. Fine-grained electrocorticography (ECoG) recordings from the middle and posterior STG reveal that phonetic features are encoded without strict spatial segregation, but rather via mixed interleaved representations [13]. At the semantic level, the anterior superior temporal gyrus (aSTG), as a part of the ATL, responds to semantic decisions from heard speech, without a clear anatomo-functional separation from the middle STG [14–18]. In functional MRI (fMRI) data, neural activity recorded through incrementally higher cognitive brain regions correlates with increasingly deeper layers in large language models [12]. These findings suggest that these brain regions represent multiple speech features, making them suitable candidates for housing transformations across the speech hierarchy.

Studies using time-resolved recording techniques such as EEG, MEG, or intracranial EEG, showed simultaneous encoding of features across the speech hierarchy, organized in increasingly larger time scales [19–21]. While phonetic features are short-lived and typically processed around 100-200 ms after phoneme onset in the STG [2, 13], semantic processing, e.g. semantic composition and lexical decisions, is associated with a longer-lasting compound event at 250 and 400 ms after word-onset [3, 22–24]. How these different time scales integrate bottom-up phonetic and top-dow semantic processes during word recognition has not yet been mechanistically characterized. A prominent model for explaining these interactions is predictive coding [25, 26]. In the context of speech processing, similar mechanisms have been operationalized in the analysis-by synthesis framework [27–30]. In these frameworks, word priors elicited by the build-up of incoming phonetic inputs are generated and directly compared to the sampled acoustic input. Depending on the amount of errors generated, the prior is either confirmed or rejected in favor of newly updated hypotheses. These frameworks thus predict that in pivotal, cross-level brain regions such as the ATL, top-down semantic predictions would meet representations of phonetic features.

To identify and characterize the fine-grained mechanisms underlying speech neural transformations, we use the framework of neural manifolds [31–38]. In the primate cortex, the dynamics in the neural manifolds distinguish different tasks and stimuli, such as sensorimotor computations, decision making, or working memory [39–41]. The invasiveness of single-neuron recordings in humans only rarely allows characterizing collective dynamics of action potentials recorded from neuronal ensembles. Yet, it seems plausible that different aspects of speech, including transformations across the speech hierarchy, are encoded in such low-dimensional manifold representations. That is, the condition- and history-dependent organization of neuronal trajectories in neural manifolds could uniquely represent the combination of phonetic and semantic features[11, 42]. Studies that have so far reported the detailed activity of populations of single units associated with speech processing have highlighted the tuning of neurons to specific speech features, leaving aside the question of the cross-level interactions [43–48]. Here, we address the mechanisms in collective neuronal dynamics underlying phonetic-to-semantic transformations.

We used unique recordings of microelectrode arrays (MEAs) implanted in the anterior STG of three patients with pharmacologically resistant focal epilepsy, who performed an auditory semantic categorization task and engaged in a spontaneous natural conversation. The MEAs were placed at the intersection of areas traditionally associated with the processing of phonetic and semantic information, while ECoG grids simultaneously recorded LFP activity along the temporal lobe. Despite the absence of detectable power effects on the most proximal ECoG channels, we observed a distributed encoding of phonemes and semantic features at the local neuronal population scale. The dynamics of individual phoneme trajectories were organized according to their corresponding phonetic features in neural manifolds. The same phonetic organization generalized to natural speech, with different speakers and a variety of complex linguistic and predictive processes. Critically, we tested whether bottom-up phonetic or top-down semantic processes organize the neuronal representations during natural speech processing. We found that phoneme encoding aligned to word onsets and concurrently with semantic features. These findings suggest that top-down semantic predictions align phonetic feature representations by predictive mechanisms, and show how phonetic representations dynamically contribute to retrieving the meaning of speech.

## Results

### Phonetic and semantic encoding in single-unit and LFP recordings of the aSTG during a semantic categorization task

To investigate the interactions between phonetic and semantic representations during speech processing, we recorded microelectrode array (MEA) and ECoG signals in three patients (Patient 1: male, 31 years old; Patient 2: male, 45 years old; and Patient 3: female, 29 years old) with pharmacologically resistant focal epilepsy, who were implanted for clinical purposes. A 10 *×* 10 MEA was positioned in the left (Participants 1 and 3) and right (Participant 2) anterior superior temporal gyrus (aSTG) (squares on Fig. 1a top and Fig. 3a top). ECoG electrodes of interest covered a large portion of the surrounding temporal cortical surface (circles on Fig. 1a top). Two of the participants (1 and 2) performed an auditory semantic categorization task (Fig. 1), and two participants (1 and 3) were additionally recorded while having a natural conversation (Fig. 3). In the auditory semantic categorization task, participants were instructed to indicate, by pressing a button, whether the heard word was smaller or larger than a foot (Fig 1a bottom). 400 unique nouns were presented, half of which indicated objects (e.g. chair), and the other half animals or body parts (e.g. donkey, eyebrow). In both groups (object, animal), words were equally divided into two categories, either larger or smaller than a foot, resulting in a balanced 2-by-2 design. To verify that phonetic and semantic processing occurred at the precise aSTG sites targeted by the intracortical MEAs, we used a large-scale fMRI database [49]. We found that MEA implantation sites are indeed involved in both phonetic processing and semantic categorization (Fig. 1b).

**Fig. 1.**
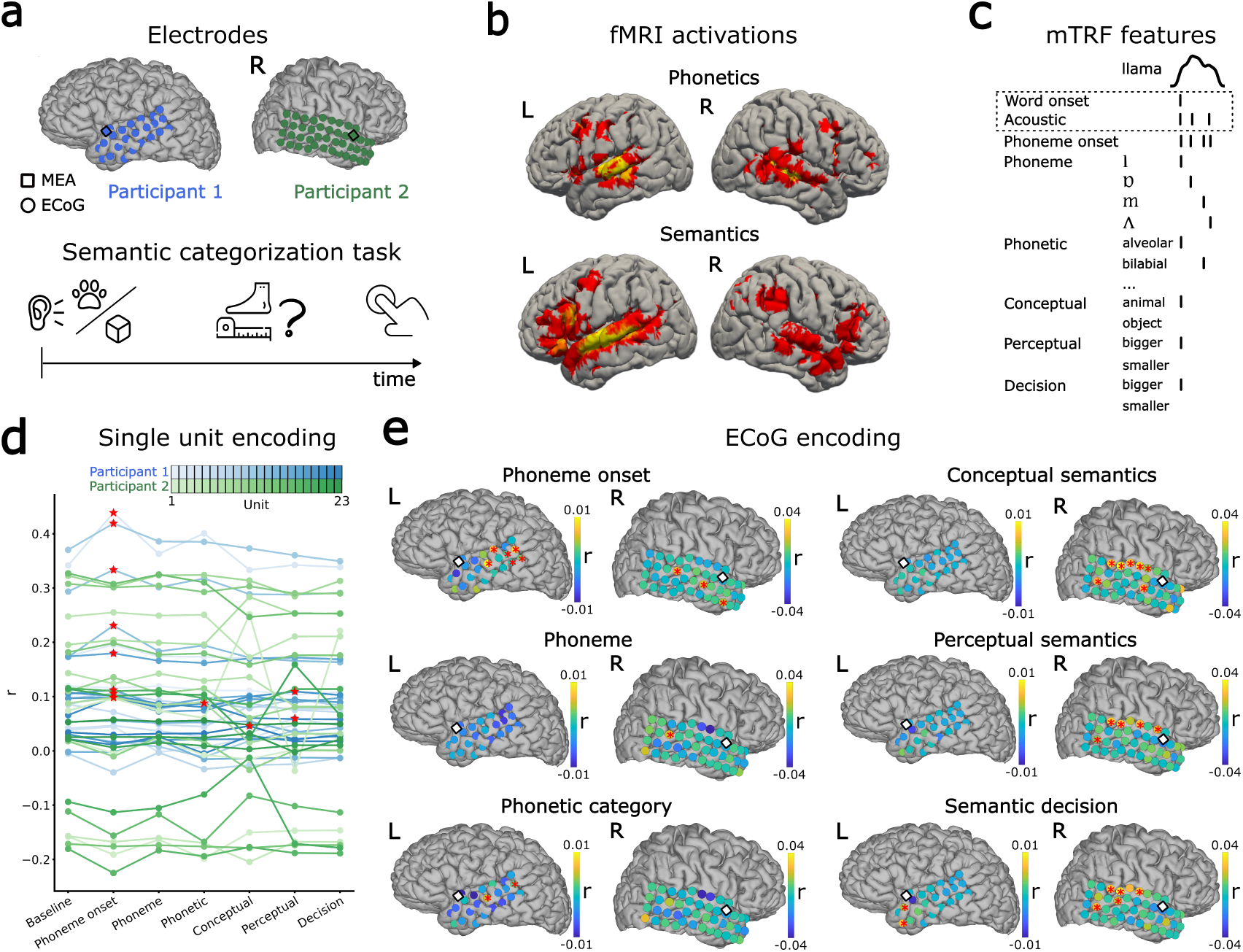
*|* **Phonetic and semantic encoding during a semantic categorization task. a.** Top: locations of the microelectrode arrays (MEA, squares) and the ECoG contacts (circles) on the cortical surface across the two participants who performed the task: Participant 1 (blue, 176 units) and Participant 2 (green, 113 units). Bottom: Auditory semantic categorization task. On each trial, participants heard a word (animal or object) and judged whether it was larger or smaller than a foot by pressing a button. **b.** fMRI correlates of phonetic processing and semantic categorization obtained from fMRI large-scale database [49]. **c.** mTRF features for the example English word ”llama”. The time series at the top indicates the speech envelope. The features used in the baseline model are indicated by a dotted square. **d.** Pearson correlation coefficient (r values) for each mTRF model fitted to each of the 23 most-spiking single units in each participant, color-coded as in a. For both participants, the lines for different units are color-coded based on firing rate, from lower (light) to higher (dark). Red stars indicate units for which r values of the fitted models are significantly higher (p *<* 0.05) than the chance-level distribution. **e** Encoding of the same mTRF models across ECoG channels. Colors indicate differences in r values compared to the baseline model. Red stars indicate significance in d. Squares indicate MEAs.

We used multivariable temporal response function (mTRF) models to contrast the encoding of different linguistic processes and speech features (see Methods, Fig. 1c), namely: (i) acoustic, including word onset and acoustic edges (envelope rate peaks) [50]; (ii) phonemic, including phoneme onset and each phoneme identity; (iii) phonetic, including features based on vowel first formant (reflecting high and low tongue positions) and second formants (front-to-back tongue position), and consonant manner of articulation (plosive, nasal, fricative, approximant, lateral approximant) and place of articulation (bilabial, labiodental, dental, alveolar, velar, uvular, and glottal); (iv) semantic, including the word’s conceptual category (object vs. animal), perceptual category (bigger or smaller than a foot), and semantic decision (participant’s response about whether the object was larger or smaller than a foot) (Supplementary tables 1, 2, and 3). The semantic decision feature was regressed separately from the perceptual category feature because the participants responded correctly in only 83.25% (Participant 1) and 82.66% (Participant 2) of the trials. Each feature of interest was added to a baseline model that included only acoustic features, and performance was compared against this baseline. Feature-specific model performance was quantified as Pearson’s correlation between the model prediction and the neuronal activity, using a 5-fold nested cross-validation procedure (Methods).

We fitted the mTRF encoding models to the soft-normalized firing rate of spike-sorted single units recorded with the MEAs (Methods) [51]. In total, we identified 289 single units across the two semantic-task participants (176 in Participant 1 and 113 in Participant 2), and selected 46 most-spiking units with an average firing rate above 0.3 spikes/second (Participant 1: mean: 1.6, sd: 1.39; Participant 2: mean: 0.37, sd: 0.06). Raster plots for these units and 80 random words are shown in the Supplementary Fig. 1a. Only a few of these units significantly responded to the probed features. For Participant 1, eight of them correlated significantly to phoneme onsets (p *<* 0.05 based on the chance level performance of a surrogate distribution, see Methods), one to phonetic features, one to conceptual categories, and two to perceptual categories (Fig. 1f). No single units responded significantly for Participant 2, possibly due to MEA’s placement in the right hemisphere.

Although several units responded significantly to different mTRF features, no features were significantly represented by the local field potentials recorded with ECoG electrodes in the immediate vicinity of the MEAs (Fig. 1e). The broadband high-frequency activity (BHA, as a proxy for local firing rate activity [52]) was assessed with equivalent mTRF models. As expected, different mTRF features were significantly encoded across different temporal-lobe channels, but no significant effects were found near MEAs. This suggests that the same aSTG location may encode different phonemic, phonetic, and semantic information at different levels of precision (firing rates vs. BHA).

Next, we examined the encoding of phonetic features within the MEA versus the most proximal ECoG electrode. To this aim, we arranged the phonetic features into four different groups (vowel first formant, vowel second formant, consonant manner of articulation, and consonant place of articulation. For each group, we contrasted a model including the corresponding phonetic features with a baseline model containing only phoneme onsets and acoustic features. For Participant 1, two units showed a significant spiking increase for consonant manner and vowel first formant models, and none for the other two models (Supplementary Fig. 2a). None of the individual phonetic groups were distinguished by the ECoG electrode adjacent to the MEAs (Supplementary Fig. 2b).

### Distributed phonetic and semantic encoding during semantic categorization task

Our previous analysis quantified the relationships between each single unit and phonemic or semantic features. Only a few single units showed significant encoding of these features. However, many units showed changes in firing rate that did not reach statistical significance when considered individually (permutation test, Fig. 1f), suggesting that encoding might emerge at the level of neuronal population dynamics. Phoneme kernels averaged across all units showed significant periods at around 200 and 400 ms in Participant 1, and around 100 ms in Participant 2, relative to the surrogate (chance-level) distribution (Fig. 2a). Considering semantic kernels, both participants showed a significant window at around 400 ms (Participant 1 for perceptual semantics and Participant 2 for semantic decision, Fig. 2a). However, only the kernels from Participant 1 around 400 ms survived multiple comparison correction (cluster-based test, see Methods). These findings indicate a mean-field population effect, motivating the hypothesis that phonetic encoding emerges from the collective dynamics of the neuronal population, which can be captured using a low-dimensional neural manifold.

**Fig. 2.**
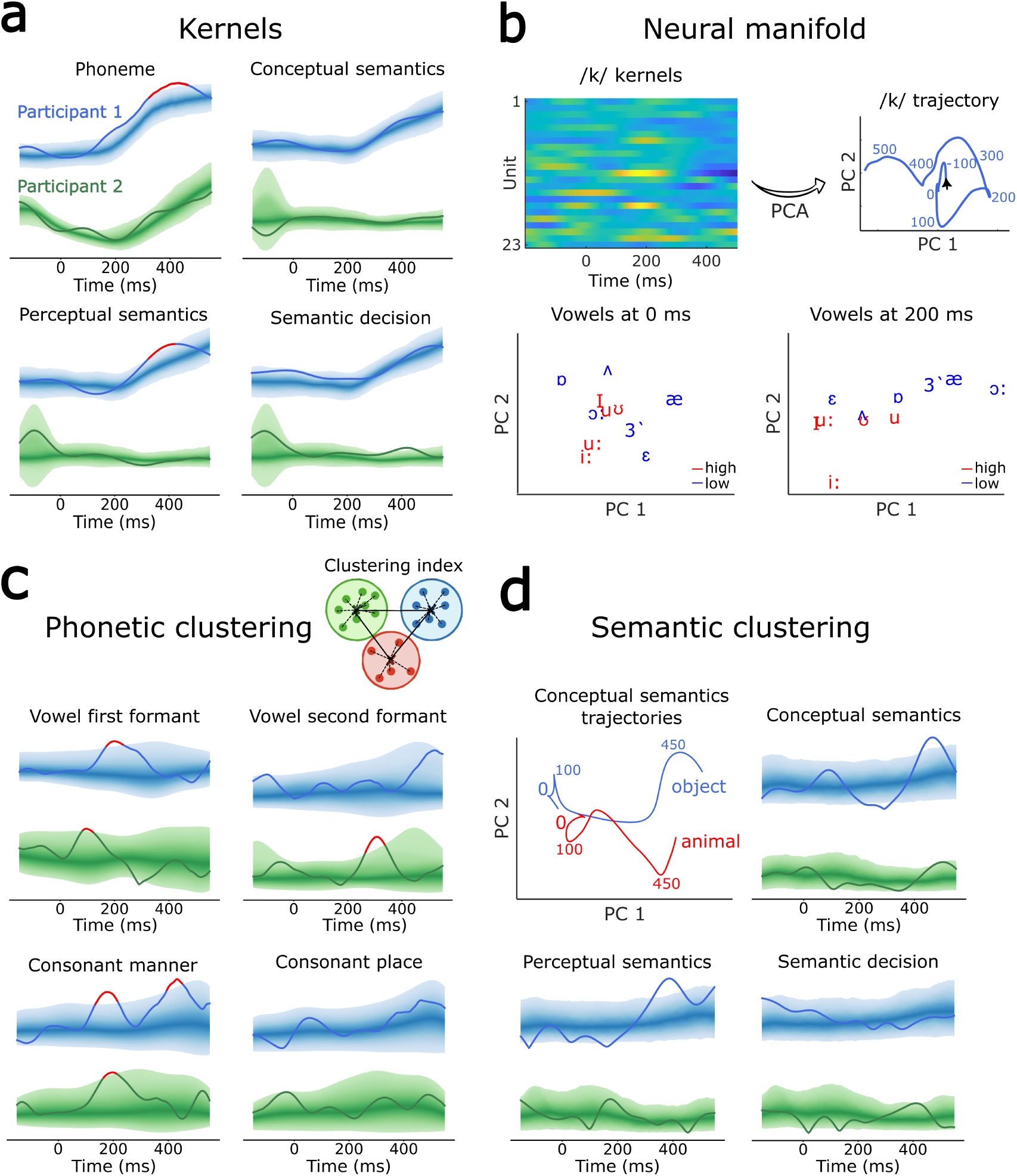
*|* **Phonetic and semantic features show distributed encoding and functional segregation in neural manifolds. a.** Average phonemic and semantic mTRF kernels for Participant 1 (blue) and Participant 2 (green). The 95% confidence region of the surrogate (chance-level) distribution is shown with brighter shading for increasingly peripheral percentiles. Red segments indicate time windows that persisted after multiple comparison correction (cluster-based test). Both participants showed several significant phonemic and semantic periods, but only Participant 1 displayed robust effects around 400 ms. **b.** Top: mTRF kernels for an example phoneme (/k/) in Participant 1, shown for the 23 most spiking units and projected into a 2D neural manifold via PCA to yield a phoneme trajectory. Bottom: Vowel distributions in a 2D manifold at 0 and 200 ms reveal a clear first-formant (tongue-height) separation at 200 ms. **c.** The clustering index—computed as the difference between between-cluster (full lines) and within-cluster (dotted lines) distances of projected phoneme trajectories—revealed formant- and articulation-based segregation of phonemes at specific time windows: vowel first-formant clustering peaked around 100–200 ms (both participants, top left), second-formant around 300 ms (Participant 2, top right), and consonant manner-of-articulation (bottom left) around 200 (both participants) and 400 ms (Participant 1), while place-of-articulation showed no significant clustering (bottom right). Coloring as in a. **d.** Top left: conceptual semantic kernels from Participant 1 projected to the two-dimensional neural manifold using PCA show a clear separation around 450 ms. Other panels: clustering indices for conceptual, perceptual, and semantic decision. In both participants, clustering of conceptual semantics occurred around 450 ms (top right). Slightly before, around 400 ms, perceptual semantic kernels separated in Participant 1 (bottom left), and semantic decision kernels in Participant 2 (bottom right). Semantic clustering effects did not persist after correcting for multiple comparisons. Coloring as in a.

**Fig. 3.**
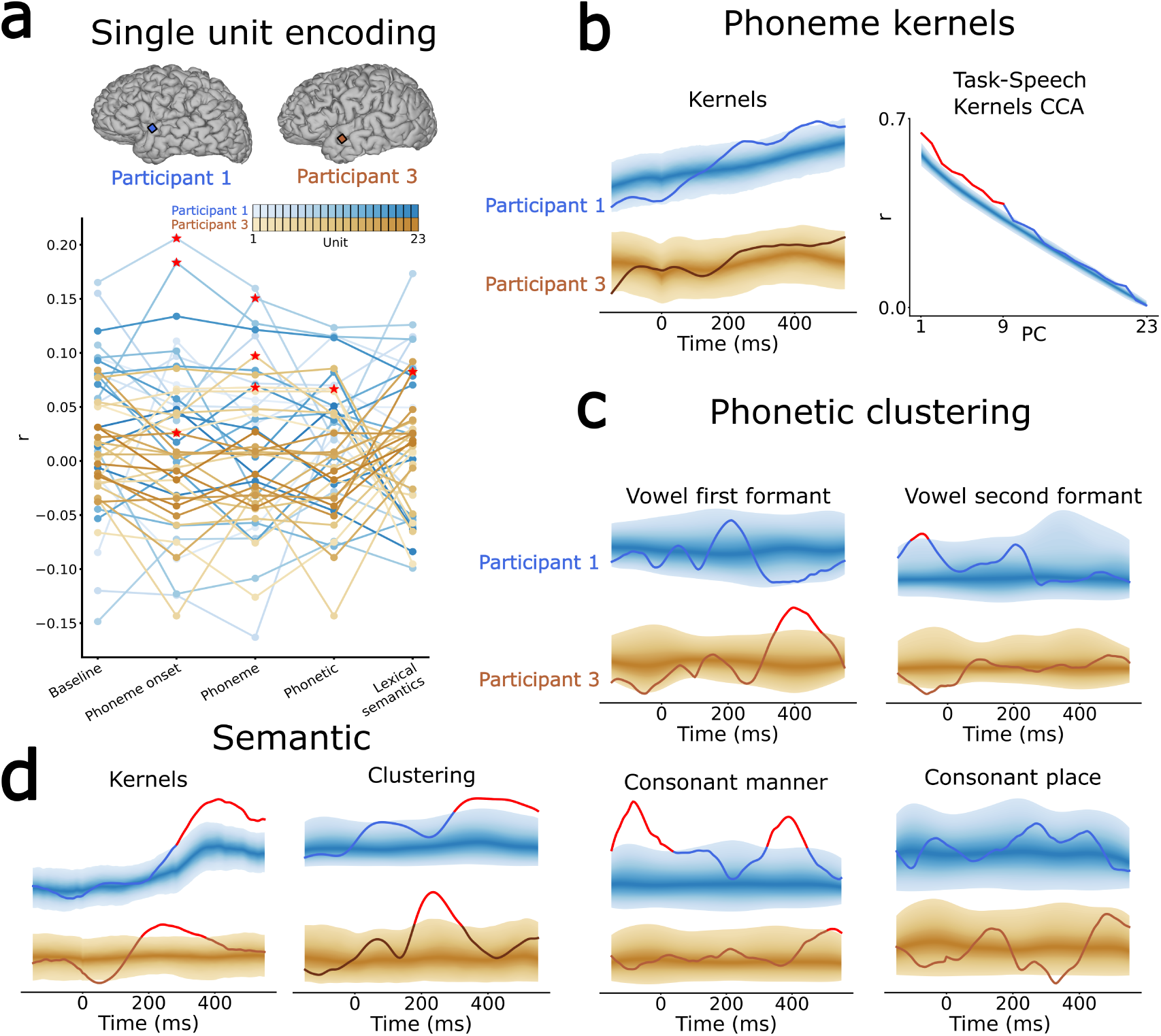
*|* **Generalization to natural speech perception a.** Top: locations of the microelectrode arrays (MEA, squares) on the cortical surface across the two participants recorded during natural speech: Participant 1 (blue, 212 units) and Participant 3 (orange, 123 units). Bottom: Pearson correlation coefficient (r values) for each mTRF model fitted to each of the 23 most-spiking single units for each participant. The lines for different units are color-coded based on firing rate, from lower (light) to higher (dark). Red stars indicate units for which r values of the fitted models are significantly higher (p *<* 0.05) than the chance-level distribution. **b.** Left: average mTRF phoneme kernels for both participants, colored as in a. The 95% confidence region of the surrogate (chance-level) distribution is shown with brighter shading for increasingly peripheral percentiles. Mirroring the semantic-task kernel effects (Fig. 2a), for Participant 1, the natural-speech kernel was significant at two time periods, centered around 200 and 400 ms. Right: Canonical correlation analysis (CCA) between the PC projections of phoneme kernels obtained during the semantic task and natural speech perception for Participant 1 showed correlations for the first nine PCs. Red segments indicate significance after multiple comparison correction. **c.** Clustering index for the four groups of phonetic features for natural-speech participants. Coloring as in b. **d.** Left: averaged lexical semantic kernels were significant at around 250 ms for Participant 3 and 400 ms for Participant 1. Right: the corresponding clustering index was significant around the same respective time periods for both participants. Coloring as in b.

To address this hypothesis, we performed principal component analysis (PCA) on all concatenated phoneme kernels obtained with mTRF for each semantic-task participant separately (Fig. 2b). The first four principal components (PCs) accounted for about 50% of phoneme feature variance for Participant 1, and about 90% for Participant 2, and were distributed across different units (Supplementary Fig. 3). Having half of the variance explained by the low-dimensional and correlated activity of several unit kernels indicates that phonemes are dynamically represented within a manifold.

We then asked whether the obtained low-dimensional neural manifold carried a functional relevance for phonetic encoding. For this, we projected all phoneme kernels onto one- to six-dimensional PC spaces and investigated whether the phonemes were grouped according to phonetic categories in the corresponding neural manifolds. Fig. 2b-top shows an example of a full trajectory for phoneme /k/ in a two-dimensional manifold for Participant 1, and Fig. 2b-bottom shows all vowel positions at 0 and 200 ms. Vowels seem to functionally segregate around 200 ms based on the first formant grouping (high vs. low tongue position). To formally quantify these spatial segregation, for each group of phonetic features (vowel first formant, vowel second formant, consonant manner, consonant place), we computed a clustering index by subtracting intra-from between-cluster distances (Fig. 2c inset) and compared it against the surrogate distribution (Methods). Mirroring the observed separation of vowels (Fig. 2b), the clustering index for vowel first formant was significant for Participant 1 around 200 ms, and, similarly, for Participant 2 around 150 ms (Fig. 2c top left). In Participant 2, vowel trajectories also separated based on the second formant (front to back tongue position) around 300 ms. In both participants, consonant trajectories clustered based on the manner of articulation (plosive, nasal, fricative, approximant, lateral approximant) around 200 ms and again around 400 ms for Participant 1 (Fig. 2c bottom left). Consonants did not segregate based on place of articulation (bilabial, labiodental, dental, alveolar, velar, uvular, and glottal features, Fig. 2c bottom right). While Fig. 2c shows clustering for two-dimensional manifolds, the same effects were observed for one-to six-dimensional manifolds (Supplementary Figs. 5 and 6). All clustering effects persisted after correcting for multiple comparisons, and they were replicated with various control analyses (Supplementary Table 5): linear discriminant analysis (LDA) (Supplementary Fig. 10), rank regression analysis (Supplementary Fig. 11), and k-means clustering (Supplementary Fig. 12).

We then turned to the semantic kernels. We projected the three semantic kernels of the semantic task (Fig. 2a) onto their corresponding low-dimensional neural manifolds (Fig. 2d and Supplementary Fig. 3), and computed the same clustering index as for phoneme trajectories. We observed a significant separation for the conceptual category kernels (object vs. animal) at about 450 ms for both participants (Fig. 2d top right), even though the averaged conceptual category kernel was never statistically significant (Fig. 2a top right). This highlights that the relevant functional read-out of certain aspects of semantic processing might emerge only at the level of the neuronal population dynamics. Slightly before the separation of conceptual semantic kernels, at around 400 ms, we also observed a significant separation of perceptual semantic kernels (bigger vs. smaller) for Participant 1 (Fig. 2d bottom left) and of semantic decision (response bigger vs. smaller) for Participant 2. The same effects persisted up to 6-dimensional PC spaces (Supplementary Figs. 7 and 8), but none survived correction for multiple comparisons.

To summarize, we observed a significant clustering of phoneme and semantic trajectories on a low-dimensional neural manifold during a semantic-categorization task, suggesting that in strategical regions involved in the integration across levels of the language hierarchy (here the aSTG), interacting speech features are jointly represented through neuronal population dynamics.

### Generalization to natural speech perception

To investigate whether the results generalized to natural speech perception, we analyzed neuronal recordings from the intracortical MEAs of two participants (1 and 3) during a spontaneous conversation with another person (Fig. 3a top), recorded in a separate experimental session. We performed spike sorting and identified 353 single units across the two natural-speech participants. To match the semantic task analysis, for each participant, we selected the 23 most-spiking units for further analysis (Participant 1: mean: 0.33 spikes/second, sd: 0.39; Participant 3: mean: 0.58 spikes/second, sd: 0.44; see Supplementary Fig. 1 for raster plots). During the speech segments, there were 664 words for Participant 1 and 1137 words for Participant 3, pronounced by a same adult person. From those words, we segmented the same 32 phonemes as in the task (Supplementary Tables 1 and 2). To investigate lexical semantics, we computed the Lancaster sensorimotor norms of each word [53] (Supplementary Table 4). We then fitted the mTRF encoding models as in the semantic task dataset, and likewise observed that only a few units significantly correlated with speech features (Fig. 3a bottom). All analyses were identical to those for the semantic task.

We first investigated the population-level effects. In Participant 1, who was recorded both during the semantic task and natural speech, we found the same significant periods in the average phoneme kernel: at about 200 and 400 ms (Fig. 3b left). While significant, these effects did not persist after controlling for multiple comparisons. In the same participant, we compared the similarity of low-dimensional phoneme trajectories obtained in the semantic task versus natural speech through canonical correlation analysis (CCA). We compared the correlations between the projected phoneme kernels of the two datasets against the surrogate distribution of canonical correlations obtained by shuffling the kernels for natural speech. We observed significant correlations for the first nine PC dimensions (Fig. 3b right), which survived multiple comparison correction. This shows that although the phonemes in the semantic task and natural speech perception are encoded by different units (as the two recordings are separated by a few hours), the neuronal population dynamics are similar, with individual phonemes tracing highly similar trajectories in the low-dimensional neural manifold.

We then proceeded as before by performing PCA on the kernels for each natural-speech participant separately. Similar to the task, 50% of the variance was explained by five PCs for both natural-speech participants, and the processing of phonemes was distributed across units (Supplementary Fig. 4). We further computed the clustering index for phonetic features in the low-dimensional neural manifold (Fig. 3c shows an example for a 2D space, and Supplementary Figs. 13 and 14 show from 1 to 6 dimensions). For Participant 1, the clustering index profiles were similar to those in the semantic task: vowels separated based on first formant around 200 ms and consonants based on manner of articulation around 400 ms. For Participant 3, vowels and consonants separated based on the same categories, but slightly later: 400 ms for vowels and 450 ms for consonants. In Participant 1, there were additional peaks at about -100 ms for vowel second formant and consonant manner of articulation, possibly reflecting prediction mechanisms present in natural speech perception but not during the semantic task. Finally, as for the semantic task, there was no separation of consonants based on place of articulation in either participant. We performed the same control analyses as for the semantic task (LDA classifier, rank regression, and k-means clustering), and replicated all the findings (Supplementary Figs. 16-18, and Supplementary Table 6).

Turning now to lexical semantics, the average semantic kernel became significant at 400 ms for Participant 1 (Fig. 3c) and at 250 ms for Participant 3, again suggesting a population effect. Clustering index in the two-dimensional neural manifold for the semantic kernels indicated separation of kernel projections around the same time periods (Fig. 3c, see Supplementary Fig. 15 for 1-to-6 dimensional manifolds).

### Top-down alignment of phonetic and semantic features

So far, we have found that both phonetics and semantics are encoded at the neuronal population level in neural manifolds, suggesting interactions between these two representations. In the predictive coding framework [25, 28], these interactions occur through hierarchical processes: bottom-up phonetic representations emanating from lower-level sensory cortices, and top-down semantic predictions coming from higher-level representations in the language hierarchy (Fig. 4a-left). However, it remains unclear how these processes align in time to allow interactions between phonetic and semantic representations. We propose two hypotheses. First, it is possible that cumulative processing of phonemic content is a pre-requisite for the semantic processing of the word (bottom-up cumulative). Second, phonemic content may be recapitulated from word comprehension (top-down alignment). If the bottom-up processes drive the alignment, phoneme representations should appear as the acoustic input unfolds, that is, at a fixed time after phoneme onset, and remain present until the word is identified (Fig. 4a-middle, hypothesis bottom-up cumulative). If, however, top-down predictive processes dictate the alignment, phoneme representations will be aligned to the top-down prediction at the word onset (Fig. 4a-right, hypothesis top-down predictive).

**Fig. 4.**
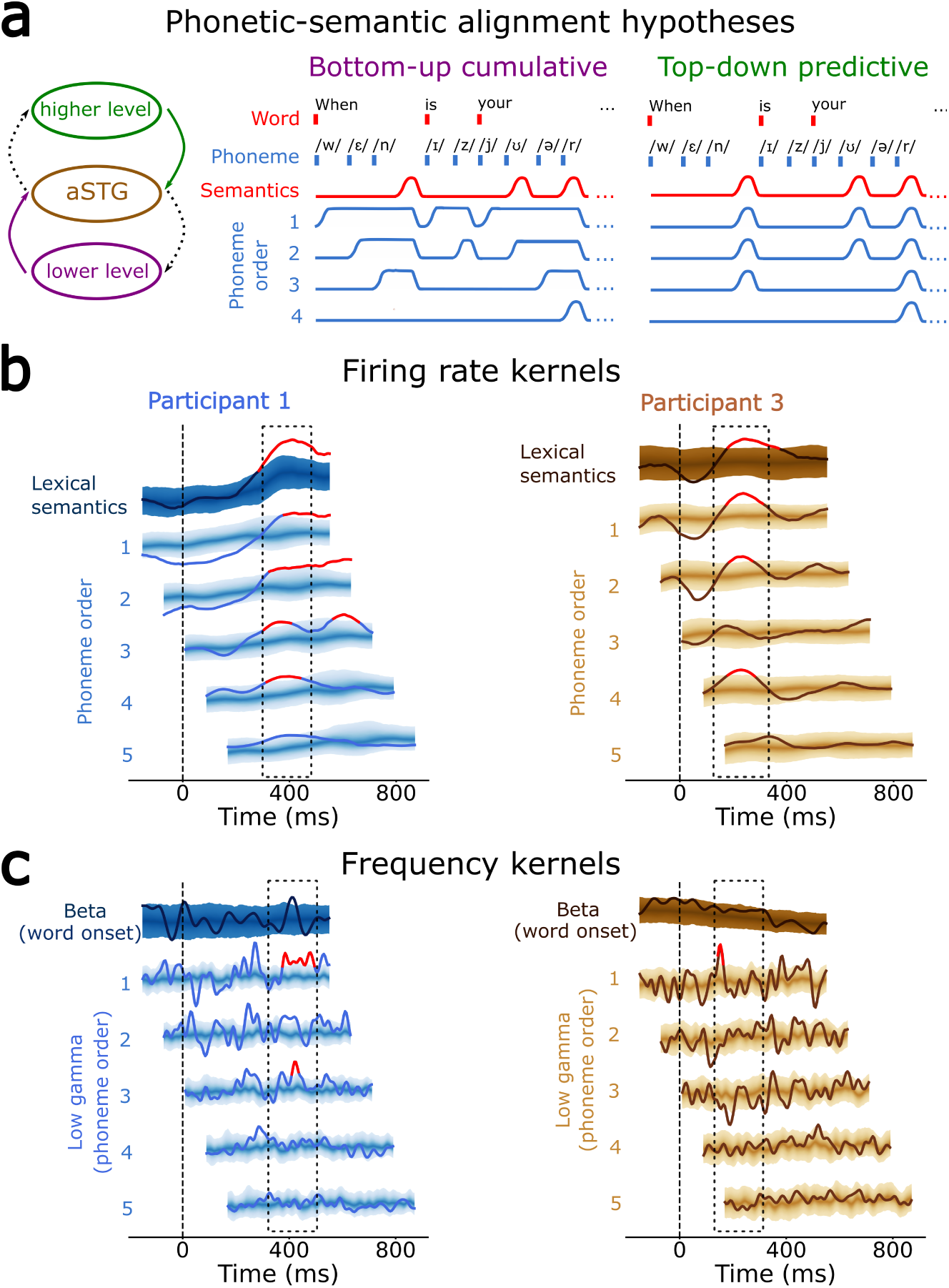
*|* **Top-down alignement of phonetic and semantic features a.** Two hypotheses about the phonetic-semantic alignment in the aSTG that might implement hierarchical processes according to the predictive coding framework (left). In the bottom-up cumulative hypothesis (middle), phoneme representations always occur at the same time after each phoneme onset as they would unfold in ongoing acoustic processes, while in the top-down predictive hypothesis (right), their sequential course disappears and they dynamically realign to the semantic representation, which occurs all at a fixed time after word onset. To test which hypothesis holds true in our data, we fitted mTRF kernels to phoneme positions within words, regardless of phoneme types (phoneme order, blue traces), and compared them to the lexical semantic kernels for each word (red traces). For example, order 2 indicates all second phonemes in all words (e.g., phoneme /E/ in ”When” and phoneme /z/ in ”is”). **b.** Phoneme order kernels aligned with the lexical semantic kernel for Participant 1 (blue) and Participant 3 (brown). For both participants, phoneme order kernels peaked simultaneously with the lexical semantic kernel at a fixed time window post-word-onset (dashed rectangle), suggesting the top-down alignment hypothesis. The 95% confidence region of the surrogate (chance-level) distribution is shown with brighter shading for increasingly peripheral percentiles. Red segments indicate significant periods after multiple comparison correction. **c.** Speech top-down processes are typically accompanied by beta peaks in the LFP power, while bottom-up processes are reflected in LFP low-gamma power peaks. We thus regressed MEA-wide beta-band activity to word onsets and low-gamma activity to phoneme orders as above. Mirroring the alignment in firing rate kernels, in Participant 1, we observed a beta-band peak at 400 ms, which co-occurred with low-gamma peaks in phoneme order kernels (dashed rectangle). There were no such frequency-band alignments in Participant 3. Coloring as in b.

To test which of the two processes drives the alignment of neural representations, we created additional mTRF features that regressed phoneme onsets at different positions within each word (i.e., phoneme order, blue traces on 4a) and compared the timing of their kernels with the timing of lexical semantic kernels (red traces on 4a). This was done by fitting an mTRF feature that regresses the order of the phonemes within the word, regardless of phoneme type. For example, order 1 is a kernel regressed from all first phonemes across all words, order 2 is a kernel regressing all second phonemes, etc. We then shifted the resulting phoneme order kernels for the corresponding multiple of the average phoneme duration (80 ms, Supplementary tables 1 and 2) - i.e., phoneme order 1 is at word onset (0 ms), order 2 at 80 ms post-word-onset, order 3 at 160 ms, etc. - and compared the alignment of the significant kernel effects. If the bottom-up cumulative hypothesis holds true, phoneme representations will start later in time as the phoneme position in the word increases, following the auditory inputs (Fig. 4a-middle); if the top-down predictive hypothesis is correct, phoneme representations will align at fixed time following the word onset, regardless of their position in the word (Fig. 4a-right). For both natural-speech participants, we observed concurrent alignment of phoneme-order kernels with lexical semantic kernels, supporting the top-down alignment hypothesis (Fig. 4b). For Participant 1, the alignment occurred at 400 ms post-word onset, while for Participant 3, this occurred at 250 ms.

To further confirm whether these alignments reflected top-down predictive or bottom-up cumulative processes, we investigated the LFP power over the MEAs. The occurrence of speech top-down processes is typically accompanied by beta peaks in the LFP power, while bottom-up processes are reflected in LFP low-gamma power peaks [54, 55]. We thus extracted and averaged the LFP power across the MEA contacts for both the beta (12-30 Hz) and low-gamma (30-70 Hz) bands. We then fitted mTRF models to these two variables by regressing word onset for the beta band (reflecting word-level top-down processing) and phoneme order for the low-gamma band (reflecting position-dependent bottom-up processing). Mirroring the lexical semantic kernel, in Participant 1, we observed a significant beta peak at 400 ms (Fig. 4c-left, top trace). Moreover, several phoneme-order peaks in the low-gamma band co-occurred with this beta peak (Fig. 4c-left, bottom traces). Some, but not all frequency-band peaks survived multiple comparison correction. For participant 3, there were no significant beta peaks co-occurring with the significant lexical semantic kernel period (Fig. 4c-right).

As an additional confirmation of the 400 ms top-down process for Participant 1, we extracted the broadband high-frequency activity (BHA, 70-150 Hz) across the MEA, as it has recently been shown to originate from both local neuronal firing and top-down predictive information [56]. We found a significant BHA-related peak at about 400 ms, coinciding with the lexical semantics peak from above (Supplementary Fig. 19a). To ensure that this peak does reflect top-down predictive information and not underlying neuronal firing [52], we also fitted a word-onset mTRF to firing rate, and observed no significant peak at 400 ms (Supplementary Fig. 19b). Finally, we investigated whether there is a causal relationship between the low-dimensional phonetic and semantic trajectories by computing Granger causality between their low-dimensional projections. We found more significant effects in the top-down direction (Supplementary Fig. 19c).

Together, these findings suggest that neuronal population dynamics in the aSTG encode both phonetic and semantic features, but are likely driven by top-down alignment processes.

## Discussion

Using auditory semantic categorization and natural speech perception in three participants with a MEA array positioned in the anterior part of the superior temporal lobe, we consistently showed that the low-dimensional neuronal population dynamics recorded during speech processing encode phonetic and semantic features simultaneously. We identified a neural manifold for both feature groups with a functional separation of their corresponding trajectories across time. Specifically, phoneme trajectories consistently clustered according to phonetic features and were highly correlated across task and natural speech conditions, suggesting invariant representations across different contexts. Similarly, semantic trajectories separated along perceptual and conceptual dimensions during the semantic task, and along the lexico-semantic dimension during natural speech. We further tested whether these phonetic and semantic processes reflected bottom-up and top-down alignment processes by examining their temporal relationships. We found that both phonetic and semantic encoding aligned to word onset during natural speech, with phonetic encoding occurring significantly earlier as the word unfolds. This suggests a top-down predictive alignemnt process, in agreement with predictive coding, and was further confirmed by the concurrent alignment of peaks in the low-gamma, high-gamma, and beta power to the word onset.

Encoding of phonetic and semantic features overlapped in space and time. The alignment of all phonemic representations with the semantic processing window (Fig. 4b), and the bidirectional causal relationship between semantic and phonemic driven activity (Supplementary Fig. 19c) advocate for predictive coding and analysis-by-synthesis frameworks, where the distinct levels are related through dynamic bottom-up and top-down predictive loops [28, 29]. This was confirmed by the synchronous transient increases in beta and low-gamma power in Participant 1 (Fig. 4c). All these representations aligned with the word-onsets rather the the phoneme-onset, indicating a top-down semantic guess that is compared to a reinstantiation of phoneme representations [57, 58]. In Participant 1, this is reminiscent of the N400 component frequently reported in the ATL, for both semantic composition and lexical decision [23, 24, 59–65], while in Participant 3, this might reflect early indication of lexical match [66] or word phonotactic probability [67, 68].

A large amount of the neuronal variance related to phonetic and semantic encoding was accounted for by low-dimensional neural manifolds. Low-dimensional phoneme trajectories clustered according to phonetic (e.g. the first formant for vowels and the manner of articulation for consonants) and semantic features (e.g. perceptual and lexical semantics), denoting a large representational flexibility. These findings experimentally support that the notion of neural manifolds also applies to neural speech processing, extending recent findings in other cognitive domains, such as sensorimotor processing, decision making, or object recognition [32, 39, 69]. The latent dynamics in the neural manifold could constitute a robust functional read-out principle for speech processing. Specifically, we observed that the encoding of speech features became more prominent when considering the coordinated dynamics of the PCs within the MEA, as opposed to the kernel activity simply averaged across units. This might account for the lack of detectable phonetic and semantic encoding at the same aSTG location in the ECoG signals, and the low-level effects for the corresponding features in our fMRI meta-analysis (Fig. 1), reflecting the average firing rate over a large population of neurons [56]. Further, for Participant 1, who was recorded both during the semantic task and natural speech, we found that the low-dimensional phoneme features traced highly correlated trajectories across the two conditions, and moreover up to the 9th PC (Fig. 4b). This is remarkable considering that the two conditions were separated by more than two hours and that the spike sorting procedure identified different units on the same array. This shows that despite the difference in the activated units, common population patterns are preserved, suggesting that the functional read-out emerges at the population dynamic level [33–35, 70].

The findings obtained in a controlled auditory semantic categorization task generalized to natural conversation for the first participant. This is particularly noteworthy, as the datasets differ in important ways. First, in the controlled task, the participants heard isolated word recordings, all of which were nouns normalized for duration and sound intensity, while focusing on a simple cognitive task – assessing the size of the heard objects and animals. Natural speech, on the other hand, involves many other cognitive and perceptual mechanisms, including connected speech that amounts to sentences containing all word types. The intrinsic difference between datasets thus allowed us to generalize the semantic findings from a carefully designed semantic categorization task to a larger category of lexical semantics. Second, even though natural speech was uttered by another speaker, we could retrieve invariant phonetic representations, both in the timing and the shape of the low-dimensional trajectories. Therefore, the low-dimensional representations we captured in the aSTG underpin speaker-normalization of phonetic representations, a notion that was suggested by LFP-level findings [71]. Third, in the natural speech task, sounds were more variable in duration and intensity than in the controlled task, entailing other processing dimensions, e.g., prosody, accents, intonation, etc. Fourth, natural speech involves a whole range of predictive processes spanning the entire speech hierarchy from low-level acoustics to syntax and semantics. Finally, our results were replicated across three participants, for which the MEAs not located in the exact same area of the aSTG. Together, these findings suggest that the low-dimensional neuronal population encoding we observed in the aSTG was robust and invariant to speech forms and contexts.

These rare human MEA recordings provide a unique opportunity to investigate neuronal population effects, at a spatial resolution that was rarely attained previously for speech, in particular dynamical effects that were not detectable even at the most adjacent ECoG contact. However, high-resolution data come with the price of a restricted spatial sampling. Our findings are specific to a small 4-by-4 mm area of aSTG encompassed by the implantation site of the MEA. This might explain why the observed phonetic effects or exact timing - e.g. vowels clustering according to the first and or the second formant - are participant-specific. It is thus possible that the aSTG is organized into small functional subdivisions and that we would observe other clustering principles based on other phonetic features if the array was implanted in nearby regions. Reciprocally we cannot exclude that similar dynamical effects could be observed if the MEA was positioned elsewhere.

To conclude, our study provides evidence for a parallel, distributed, and low-dimensional encoding of phonetic and semantic features that is specific to neuronal population firing patterns in a focal region in the aSTG. These local dynamic population effects are part of bottom-up and top-down dynamics involving oscillatory bidirectional activity potentially involving higher- and lower-tier regions of the language hierarchy. Extending the rapidly emerging neural manifold framework to speech processing, these findings shed new light on the brain mechanisms underlying phonetic and semantic integration and pave the way toward the elucidation of the intricacies behind the complex transformations across the speech processing hierarchy.

## Methods

### Participants

The study involved three participants with pharmacologically resistant epilepsy (focal seizures), who underwent intracranial electrode implantation for neuromonitoring of epileptogenic brain areas as part of their clinical epilepsy treatment. At the time of the experimental recordings, Participant 1 was a 31-year-old male; Participant 2 was a 45-year-old male; and Participant 3 was a 29-year-old female. They were native English speakers with normal sensory and cognitive functions and demonstrated left-hemisphere language dominance through a WADA test.

Informed consent was obtained for all participants, and the study was conducted under the oversight of the Massachusetts General Hospital Institutional Review Board (IRB). The study included both the intracortical implantation of a microelectrode array (MEA) and the performance of a semantic task and natural speech. The MEA recordings were used only for scientific research purposes and played no role in the clinical assessments and decisions.

### Neural recordings

Intracranial cortical field potentials were recorded with subdural ECoG arrays (Adtech Medical), 1-cm electrode distance. In this study, only ECoG recordings from Participants 1 and 2 were examined. Specifically, electrodes covering the lateral temporal lobe were included in the analysis (Fig 1a). The signal was recorded with a sampling rate of 500 Hz, with a bandpass filter spanning 0.1 to 200 Hz. All electrode positions were accurately localized relative to the participants’ reconstructed cortical surface [72].

Single-unit action potentials were recorded with a 10-by-10, 400 µm electrode distance, microelectrode array (Utah array, Blackrock Neurotech) surgically implanted within the left anterior superior temporal gyrus (aSTG) for Participants 1 and 3 (Fig 1a), and the right aSTG for Participant 2 (Fig 3a). Electrodes were 1.5 mm long and contained a 20-µm platinum tip. The implantation sites were excised, and the subsequent histological analysis revealed the spatial orientation of the electrode tips within the depths of cortical layer III with no notable histological abnormalities in the neighboring cortical environment. Data acquisition was acquired with a Blackrock NeuroPort system, with a sampling frequency of 30 kilosamples per second, and an analog bandpass filter ranging from 0.3 Hz to 7.5 kHz for antialiasing.

The location of the recording arrays was based on clinical considerations. In particular, the MEA were placed in the superior temporal gyrus because this was a region within a larger area anticipated to be resected based on prior imaging data.

### Auditory stimuli

In Participant 1, neural data were recorded during two separate experimental sessions, which took place on the same day, two hours apart [43]. In the first session, the participant performed an auditory semantic categorization task. The stimuli were standalone audio files of 400 words pronounced by a male speaker and normalized for intensity and duration (500 ms). The participant was presented with 800 nouns in a randomized order, and with 2.2 s stimulus onset asynchrony. Out of 800 words, 400 were presented only once, while the remaining 400 consisted of 10 words repeated 40 times each. To avoid biasing effects, in our analysis, we considered only the 400 words that were repeated only once. Specifically, the inclusion of repeated words leads to the over-representation of a few phonemes compared to other phonemes, biasing the regression analyses. Half of the 400 unique words referred to objects and half to animals. Following a word presentation, the participant was instructed to press a button if the referred item was bigger than a foot in size. Half of the items in each group (animals, objects) were bigger than a foot, resulting in a balanced 2-by-2 design.

In the second session, the participant engaged in a conversation with another person present in the room. The natural speech was recorded using a far-field microphone and manually transcribed. The recording was split into 91 segments that contained clear speech recordings of the other person talking (i.e. without overlapping speech or other background sounds). Each segment was cleaned for background noise and amplified to 0 dBFS in Audacity software. We used a total of 664 words (272 unique) across all trials of the natural speech (Supplementary Table 4).

In Participant 2, the procedure was identical to the first session of Participant 1 (the semantic task). In Participant 3, a natural speech segment similar to the second session of Participant 1 was used. The recording was split into 178 segments with clear speech, and we used a total of 1137 words (431 unique).

### Signal preprocessing

Spike detection and sorting were performed with the semi-automatic wave clus algorithm [73]. In Participant 1, across 96 active electrodes, we identified 176 and 212 distinct units for the sessions with the semantic task and the natural speech, respectively. For the semantic task, we considered units with firing rate higher than 0.3 spikes/second, resulting in 23 units (mean: 1.6, sd: 1.39 spikes/s). For a fair comparison across both sessions, we also selected the 23 most spiking units in the natural speech (mean: 0.33, sd: 0.39 spikes/s). In Participant 2, we identified 113 units (mean: 0.37, sd: 0.06 spikes/s), and in Participant 3, 123 units (mean: 0.58, sd: 0.44 spikes/s). Raster plots for all participants are shown on Supplementary Fig. 1.

The spike train of each unit was smoothed with a 25 ms wide Gaussian kernel to obtain the firing rate time series. Firing rate time series were then soft-normalized by the range of the unit increased with a constant 5, as performend in previous studies, so that neurons with very high-firing rates remain close to the unity range [51]. Firing rate time series were then downsampled at 200 Hz. For the semantic task, firing rate time series were split into 400 trials. Each trial lasted 1.5 seconds and included 0.5-second periods before and after the word presentation. For the natural speech, we selected 91 segments (Participant 1) and 178 segments (Participant 3) of the firing rate where a person was talking to the participant (see above).

The signals from the ECoG grids were first filtered to remove line noise using a notch filter at 60 Hz and harmonics (120, 180, and 240 Hz). We then applied common-average referencing. For each channel, we extracted broadband high-frequency activity (BHA) in the 70-150 Hz range [52]. BHA was computed as the average z-scored amplitude of eight band-pass Gaussian filters with center frequencies and bandwidth increasing logarithmically and semi-logarithmically respectively. The resulting BHA was downsampled to 100 Hz.

#### Phoneme segmentation and categorization

Audio files and corresponding transcripts were segmented both into words and phonemes by creating PRAAT TextGrid files [74] through WebMAUS software [75]. Phonetic symbols in the resulting files were encoded in X-SAMPA, a phonetic alphabet designed to cover the entire range of characters in the 1993 version of the International Phonetic Alphabet (IPA) in a computer-readable format. All TextGrids were manually inspected and converted into tabular formats using the TEICONVERT tool. Diphtongues and phonemes that occurred less than 5 times throughout the entire session were removed from the analysis. We used 32 segmented phonemes, divided into 11 vowels and 21 consonants, and further labeled according to the standard IPA phonetic categorizations (Supplementary Tables 1 and 2): vowels first formant (high, low), vowels second formant (front, back), consonants articulation place (bilabial, labiodental, alveolar, velar, uvular, glottal), consonants articulation manner (plosive, nasal, fricative, approximant, lateral approximant).

### mTRF features

For both conditions (semantic task and natural speech), we extracted the following features: word onset, acoustic edge (envelope rate peaks), phoneme onset, phoneme identity, and phonetic category. For the semantic task, we additionally computed the following semantic features: perceptual category, conceptual category, and semantic decision. For the natural speech, we additionally created lexical semantics feature. All features were designed as values located at the onset of the corresponding stimuli, all other values being set to zeros.

Word onset was marked by a value of one located at the onset of each word. Acoustic edges were defined as local maxima in the derivative of the speech envelope [50]. The speech envelope was computed as the logarithmically scaled root mean square of the audio signal using the MATLAB mTRFenvelope function. Phoneme onset feature indicates onsets of all phonemes in a word. Phoneme identity feature was multivariable with 32 regressors, each indicating onsets of a different phoneme, as defined by the IPA table. Phonetic category feature was multivariable, and included four phonetic groups (vowel first and second formant, consonant manner, and place of articulations). All other features were multivariables with a value located at the corresponding word onset. Perceptual category, conceptual category, and semantic decision had two regressors, defined respectively as bigger and smaller, animal and object, and decision on whether the object/animal was bigger or smaller than a foot. Finally, the lexical semantics feature was multivariable, designed by regressing each of the 11 sensorimotor norms at the corresponding word onset [53]. All stimuli were smoothed with a 25-ms-wide Gaussian kernel and downsampled to either 200 Hz (to match single unit firing rates) or 100 Hz (to match BHA from ECoG channels) before fitting the mTRF models.

### mTRF estimation

mTRFs were estimated using the mTRF MATLAB toolbox [76]. All mTRFs were always of encoding type, relating the stimulus features to neural data, with resulting kernels in the time range between -200 and 600 ms. The first and last 50 ms were not considered in the analysis, to avoid possible edge effects. Both for semantic task and natural speech, the baseline model contained the word onset and the acoustic onset edge features. All other models included the baseline features and one of the additional target features defined above. Estimation was performed by a ridge regression, using a nested cross-validation procedure (see below). The goodness of fit was defined as Pearson’s correlation between model prediction and neural data.

#### Cross-validation

For the semantic task, we performed a nested cross-validation. In the outer cross-validation loop, we split the 400 words randomly into 5 sets of 80 (20% of the words each). Thus, in each fold, 80 trials belonged to a hold-out test set, while the remaining 320 words belonged to the train set. The five folds were identical across all models. For a given outer fold, among the 320 train-set trials, we performed another 8-fold inner cross-validation loop for the ridge regression hyperparameter tuning, with *λ* ranging from 10*^−^*^6^ to 10^6^. The optimal lambda was then used to retrain the model on the 320 words of the training set of the outer cross-validation loop fold. The model predictions and Pearson’s correlation with the neural data were then computed for the 80 words of the test set. Across the 5 folds, we thus obtained five correlation values, of which we report the average.

For natural speech, we also used nested cross-validation. The 5-fold outer cross-validation loop is performed by splitting the speech segments into five folds of approximately similar duration (mean: 393.28 s; sd: 6.73 s), chosen through random shuffling across the five folds until the standard deviation of the duration distribution across folds was smaller than 10 seconds. We applied a similar procedure for the inner cross-validation loop in each of the five folds.

#### Surrogate distributions and statistical significance

For each model of the semantic task, we created a distribution of 1000 surrogate models by shuffling the target feature across words, and keeping the baseline features constant. For instance, in the model that contained the phoneme onset feature together with the baseline features (word onset and acoustic edges), the surrogate model was created by randomly shuffling the 400 phoneme onset features across 400 words independently, while keeping the order of the baseline features constant. In this way, the baseline features were properly regressed to the neural data, while the target feature (e.g. phoneme onset) was randomly assigned to neural data.

For the natural speech condition, it was not possible to shuffle trials in the same way, as each auditory segment was of a different duration. This posed a problem because the mTRF features have to be of the same length as the neural data, which was not the case for natural speech, contrary to the semantic task where each word had the same duration. Instead, we used the following method: for multivariable features, the surrogates were computed by randomly assigning each non-zero value to a particular regressor (e.g. for the phoneme identity feature, the first phoneme is randomly assigned to any of the 32 regressors, the second to any of the remaining 31, etc). For features with a single variable, surrogates were computed by performing a circular shift with a random onset. For instance, for the phoneme onset feature, which had only one regressor, we randomly split the trace into 2 parts and switched the order of the parts.

A model was considered statistically significant if the original model performed better than the 95th percentile of the surrogate distribution (one-tailed test).

### Averaged kernels

To obtain averaged kernels, we took the mean of kernels across all units (and phonemes for the phoneme kernels). An averaged kernel was considered statistically significant if higher or lower than the 97.5th or 2.5th percentile respectively of the surrogate distribution (two-tailed test).

#### Correcting for multiple comparisons

To control for multiple comparisons, we used a nonparametric approach based on the cluster mass test [77]. From each permutation, we subtracted the mean value of the surrogate distribution and clustered consecutive time points that were outside the 95% confidence region of the surrogate distribution. Cluster-level statistics was defined as the sum of the mean-centered values within a cluster. For negative clusters, we considered the absolute value of the sum. Clusters of the observed data that were higher than the 95th percentile of the distribution of maximum cluster surrogates were considered as passing the multiple comparison correction.

### Clustering of phoneme kernels in the PC space

Clustering was performed by assigning each phoneme to a particular phonetic class and computing the clustering index, defined as the difference between between-cluster and intra-cluster distances. Specifically, for each cluster, we first found the location of the centroid by averaging the coordinates of all cluster elements. Between-cluster distance was defined as the average Euclidean distance between all pairs of cluster centroids. Intra-cluster distance was defined as the Euclidean distance of each cluster element to the corresponding centroid. By subtracting intra- from between-cluster distance, our index rewarded cluster separability (between-cluster distance) and penalized spatial dispersion (intra-cluster distance).

We performed clustering in the two-dimensional PC space (Results), but systematically confirmed our results in the PC spaces using up to six dimensions (Supplementary Material). Clustering was considered statistically significant if the original model had a higher clustering index than the 95th percentile of the surrogate distribution (one-tailed test).

To confirm our clustering results, we performed several control analyses, described here shortly and in detail in Supplementary Material:

1. linear discriminant analysis (LDA) classifier: at each time point and for each phonetic feature group, we first ran an LDA classifier to compute the means of the multivariate normal distributions of phonemes sharing the same phonetic feature. Then, we computed the average Euclidean distance between all phonetic feature means and compared this distance against the distribution of 1000 surrogates.
2. rank regression for vowels: we additionally explored whether the actual first and second formant frequency values were encoded in the low-dimensional space. To that aim, we assigned a rank value (1-7) to each vowel, based on the formant values indicated in the standard IPA table. At each time point, the ranked order of vowels was correlated with their coordinates on the first three PCs, and compared against a distribution of 1000 surrogates.
3. correlation with K-means connectivity matrices: to investigate whether the same results would emerge in a data-driven fashion, for each time point, we ran K-means clustering 1000 times (clustering results slightly differed depending on the algorithm’s random initialization). Then, we computed an average N-by-N connectivity matrix that indicated how often each of the N phonemes was clustered together. Finally, we correlated the resulting connectivity matrix with the connectivity matrix of the actual, phonetic-based clusters (vowel first formant, vowel second formant, consonants manner, consonants place), and compared the correlation value against the distribution of 1000 surrogates.

### Clustering of semantic feature kernels in the PC space

Each semantic feature of the semantic task only had two regressors (e.g. animal vs object, larger vs smaller), hence the same clustering algorithm was reduced to a between-cluster distance (intra-cluster distance was zero, as there was only one element in each cluster). To identify the periods during which the two kernels of each semantic feature were significantly separated in the PC space, we computed the average Euclidean distance between the two. The same process was repeated for each of the 1000 surrogate models, and the distance was considered significant when higher than the 95th percentile of the surrogate distribution (one-tailed test).

### Comparison of trajectories between the semantic task and natural speech

In Participant 1, who was recorded during both conditions (the task and natural speech), we investigated the similarity of the trajectories of the phoneme kernels projected to the low-dimensional space across the two conditions. We computed the canonical correlation between each kernel pair (e.g. kernel of phoneme /k/ extracted during the semantic task and the /k/ kernel from the natural speech). The canonical correlation was compared against the distribution of surrogate model canonical correlations for all 23 dimensions. Particularly, the surrogate distribution was computed by shuffling the kernels in the natural speech, which constitutes a strong surrogate test. The correlation was considered significant if it was higher than the value of the 95th percentile of the surrogate correlation distribution (one-tailed test).

### Phoneme order kernels

To probe for interactions between the encoding of phonetic and semantic features during natural speech, we first created mTRF kernels for phoneme onsets at each phoneme position within the word and then aligned the resulting kernels with the corresponding semantics kernel. Phoneme order was thus a multivariable feature with five regressors, each indicating the onset of the corresponding phoneme position across all words. For instance, phoneme order 2 indicates the onset of second phonemes in all words, regardless of the phoneme type. The resulting kernels were then shifted by the multiples of 80 ms, which is a rounded value of average phoneme duration (mean: 82.6 ms, sd: 3.1 ms, Supplementary Tables 1 and 2). Thus, the kernel for position two was shifted by 80 ms, the kernel for position three by 160 ms, etc. Surrogate kernel distributions were computed as described above. Phoneme onset kernels are considered significant if they are higher or lower than the value of the 97.5th or 2.5th percentile respectively of their surrogate kernel distributions (two-tailed test). This allowed us to observe when significant peaks for each phoneme occurred in reference to the same time point: word onset.

### LFP beta and low-gamma power analysis

We further investigated the nature of these kernels within the analysis-by-synthesis framework, by fitting mTRF encoding-type models with either word onset or position-based phoneme onset as stimulus, and either beta (12 - 30 Hz) or low-gamma (30 - 70 Hz) LFP power as response. Word onset and phoneme onset stimuli were the same as the ones used before. For each microelectrode channel, LFP power bands were computed by first applying a 9-order bandpass Butterworth filter with zero-phase forward and reverse digital filtering, then subtracting the mean from the resulting trace, and finally computing the absolute value of its Hilbert transform. The resulting LFP powers were then averaged across all channels and entered into the mTRF models. Finally, we compared the significant periods of the word-onset beta-power kernel and phoneme-order low-gamma kernels with the same aligning procedure as above. In the same way, we computed the BHA (70 - 150 Hz) word onset kernel, and compared it to the firing rate word onset kernel (Supplementary Figure 19).

### Granger causality

We adopted a Granger causality measure [78] to explore the temporal causality between the low-dimensional phonetic and semantic representations. We used the multivariate Granger causality toolbox (MVGC v1.3, 2022), which is based on a state-space formulation of Granger causal analysis [79, 80]. We first applied a half-Gaussian filter 25 ms wide to the spiking traces of individual units to obtain a causal firing rate estimate. The resulting firing rates were then projected into the three-dimensional phonetic and semantic PC spaces, constructed by performing PCA on the corresponding phonetic and semantic mTRF kernels as described above. The Granger models were created by separately predicting each of the three PCs of one feature (e.g. phonetic) from all three dimensions of another feature (e.g. semantic). We then combined the resulting Granger coefficients into two 3-by-3 matrices, one for phonetic-to-semantic and another one for semantic-to-phonetic predictions (Supplementary Figure 19c). Autoregression model parameters were estimated from data via the Levinson-Wiggins-Robinson algorithm. The order of AR models was selected via the Akaike Information Criterion (AIC). Statistical significance of estimated Granger causality measures was assessed via the corresponding F-test as implemented by the toolbox.

#### fMRI databases

Functional classification of the cortical surface surrounding the MEA with respect to different linguistic processes was performed using NeuroSynth Compose, a platform for neuroimaging meta-analyses. We were primarily interested in observing the proximity of phonetic and semantic processing close to the MEA implantation site. Relevant studies were found with keyword and then manually curated. The full meta-analyses are available in neurovault for both the phonetic (https://identifiers.org/neurovault.collection:22048) and semantic (https://neurovault.org/collections/22048/) meta-analyses.

## Acknowledgments

This work was funded by Swiss National Science Foundation career grant 193542 and 225979 (T.P.), EU FET-BrainCom project (A.G.), NCCR Evolving Language, Swiss National Science Foundation Agreement #51NF40 180888 (A.G.), Swiss National Science Foundation project grant 163040 (A.G.), National Institutes of Health (NIH), National Institute of Neurological Disorders and Stroke (NINDS), grants R01NS079533 (WT) and R01NS062092 (SSC), and the Pablo J. Salame Goldman Sachs endowed Associate Professorship of Computational Neuroscience at Brown University (WT).

## References

[1] Hickok, G. & Poeppel, D. The cortical organization of speech processing. Nature Reviews Neuroscience 8, 393–402 (2007).

[2] Mesgarani, N., Cheung, C., Johnson, K. & Chang, E. F. Phonetic Feature Encoding in Human Superior Temporal Gyrus. Science 343, 1006–1010 (2014).

[3] Pylkkanen, L. Neural basis of basic composition: What we have learned from the red–boat studies and their extensions. Philosophical Transactions of the Royal Society B: Biological Sciences 375, 20190299 (2020).

[4] Patterson, K., Nestor, P. J. & Rogers, T. T. Where do you know what you know? The representation of semantic knowledge in the human brain. Nature Reviews Neuroscience 8, 976–987 (2007).

[5] Ralph, M. A. L., Jefferies, E., Patterson, K. & Rogers, T. T. The neural and computational bases of semantic cognition. Nature Reviews Neuroscience 18, 42–55 (2017).

[6] Noppeney, U. et al. Temporal lobe lesions and semantic impairment: A comparison of herpes simplex virus encephalitis and semantic dementia. Brain 130, 1138–1147 (2006).

[7] Lambon Ralph, M. A., Lowe, C. & Rogers, T. T. Neural basis of category-specific semantic deficits for living things: Evidence from semantic dementia, HSVE and a neural network model. Brain 130, 1127–1137 (2006).

[8] Schwartz, M. F. et al. Anterior temporal involvement in semantic word retrieval: Voxel-based lesion-symptom mapping evidence from aphasia. Brain 132, 3411–3427 (2009).

[9] Cope, T. E. et al. Anterior temporal lobe is necessary for efficient lateralised processing of spoken word identity. Cortex 126, 107–118 (2020).

[10] de Heer, W. A., Huth, A. G., Griffiths, T. L., Gallant, J. L. & Theunissen, F. E. The Hierarchical Cortical Organization of Human Speech Processing. The Journal of Neuroscience 37, 6539–6557 (2017).

[11] Yi, H. G., Leonard, M. K. & Chang, E. F. The Encoding of Speech Sounds in the Superior Temporal Gyrus. Neuron 102, 1096–1110 (2019).

[12] Caucheteux, C., Gramfort, A. & King, J.-R. Deep language algorithms predict semantic comprehension from brain activity. Scientific Reports 12, 16327 (2022).

[13] Hamilton, L. S., Oganian, Y., Hall, J. & Chang, E. F. Parallel and distributed encoding of speech across human auditory cortex. Cell 184, 4626–4639.e13 (2021).

[14] Scott, S. K. Identification of a pathway for intelligible speech in the left temporal lobe. Brain 123, 2400–2406 (2000).

[15] Visser, M. & Lambon Ralph, M. A. Differential Contributions of Bilateral Ventral Anterior Temporal Lobe and Left Anterior Superior Temporal Gyrus to Semantic Processes. Journal of Cognitive Neuroscience 23, 3121–3131 (2011).

[16] Chang, E. F., Raygor, K. P. & Berger, M. S. Contemporary model of language organization: An overview for neurosurgeons. Journal of Neurosurgery 122, 250–261 (2015).

[17] Zhang, Y. et al. Hierarchical cortical networks of “voice patches” for processing voices in human brain. Proceedings of the National Academy of Sciences 118, e2113887118 (2021).

[18] Damera, S. R. et al. Evidence for a Spoken Word Lexicon in the Auditory Ventral Stream. Neurobiology of Language 4, 420–434 (2023).

[19] Heilbron, M., Armeni, K., Schoffelen, J.-M., Hagoort, P. & De Lange, F. P. A hierarchy of linguistic predictions during natural language comprehension. Proceedings of the National Academy of Sciences 119, e2201968119 (2022).

[20] Gwilliams, L., Marantz, A., Poeppel, D. & King, J.-R. Hierarchical dynamic coding coordinates speech comprehension in the brain (2024).

[21] Keshishian, M. et al. Joint, distributed and hierarchically organized encoding of linguistic features in the human auditory cortex. Nature Human Behaviour 7, 740–753 (2023).

[22] Friederici, A. D. & Kotz, S. A. The brain basis of syntactic processes: Functional imaging and lesion studies. NeuroImage 20, S8–S17 (2003).

[23] Kutas, M. & Federmeier, K. D. Thirty Years and Counting: Finding Meaning in the N400 Component of the Event-Related Brain Potential (ERP). Annual Review of Psychology 62, 621–647 (2011).

[24] Dikker, S., Assaneo, M. F., Gwilliams, L., Wang, L. & Kösem, A. Magnetoencephalography and Language. Neuroimaging Clinics of North America 30, 229–238 (2020).

[25] Rao, R. P. & Ballard, D. H. Predictive coding in the visual cortex: A functional interpretation of some extra-classical receptive-field effects. Nature neuroscience 2 (1999).

[26] Friston, K. & Kiebel, S. Predictive coding under the free-energy principle. Philosophical Transactions of the Royal Society B: Biological Sciences 364, 1211–1221 (2009).

[27] Mackay, D. M. Towards an information-flow model of human behavior. British Journal of Psychology 47, 30–43 (1956).

[28] Halle, M. & Stevens, K. Analysis by synthesis. Proceeding of the Seminar on Speech Compression and Processing **II**, 1 (1959).

[29] Bever, T. G. & Poeppel, D. Analysis by Synthesis: A (Re-)Emerging Program of Research for Language and Vision. Biolinguistics 4, 174–200 (2010).

[30] Su, Y., MacGregor, L. J., Olasagasti, I. & Giraud, A.-L. A deep hierarchy of predictions enables online meaning extraction in a computational model of human speech comprehension. PLOS Biology 21, e3002046 (2023).

[31] Pillai, A. S. & Jirsa, V. K. Symmetry Breaking in Space-Time Hierarchies Shapes Brain Dynamics and Behavior. Neuron 94, 1010–1026 (2017).

[32] Gallego, J. A., Perich, M. G., Miller, L. E. & Solla, S. A. Neural Manifolds for the Control of Movement. Neuron 94, 978–984 (2017).

[33] Jazayeri, M. & Ostojic, S. Interpreting neural computations by examining intrinsic and embedding dimensionality of neural activity. Current Opinion in Neurobiology 70, 113–120 (2021).

[34] Vyas, S., Golub, M. D., Sussillo, D. & Shenoy, K. V. Computation Through Neural Population Dynamics. Annual Review of Neuroscience 43, 249–275 (2020).

[35] Chung, S. & Abbott, L. Neural population geometry: An approach for understanding biological and artificial neural networks. Current Opinion in Neurobiology 70, 137–144 (2021).

[36] Martin, A. E. A Compositional Neural Architecture for Language. Journal of Cognitive Neuroscience 32, 1407–1427 (2020).

[37] Truccolo, W. From point process observations to collective neural dynamics: Nonlinear Hawkes process GLMs, low-dimensional dynamics and coarse graining. Journal of Physiology-Paris 110, 336–347 (2016).

[38] Aghagolzadeh, M. & Truccolo, W. Inference and Decoding of Motor Cortex Low-Dimensional Dynamics via Latent State-Space Models. IEEE Transactions on Neural Systems and Rehabilitation Engineering 24, 272–282 (2016).

[39] Mante, V., Sussillo, D., Shenoy, K. V. & Newsome, W. T. Context-dependent computation by recurrent dynamics in prefrontal cortex. Nature 503, 78–84 (2013).

[40] Remington, E. D., Narain, D., Hosseini, E. A. & Jazayeri, M. Flexible Sensorimotor Computations through Rapid Reconfiguration of Cortical Dynamics. Neuron 98, 1005–1019.e5 (2018).

[41] Markowitz, D. A., Curtis, C. E. & Pesaran, B. Multiple component networks support working memory in prefrontal cortex. Proceedings of the National Academy of Sciences 112, 11084–11089 (2015).

[42] Pulvermüller, F. Neural reuse of action perception circuits for language, concepts and communication. Progress in Neurobiology 160, 1–44 (2018).

[43] Chan, A. M. et al. Speech-Specific Tuning of Neurons in Human Superior Temporal Gyrus. Cerebral Cortex 24, 2679–2693 (2014).

[44] Chan, A. M. et al. First-Pass Selectivity for Semantic Categories in Human Anteroventral Temporal Lobe. Journal of Neuroscience 31, 18119–18129 (2011).

[45] Ossmy, O., Fried, I. & Mukamel, R. Decoding speech perception from single cell activity in humans. NeuroImage 117, 151–159 (2015).

[46] Lakretz, Y., Ossmy, O., Friedmann, N., Mukamel, R. & Fried, I. Single-cell activity in human STG during perception of phonemes is organized according to manner of articulation. NeuroImage 226, 117499 (2021).

[47] Leonard, M. K. et al. Large-scale single-neuron speech sound encoding across the depth of human cortex. Nature 1–10 (2023).

[48] Khanna, A. R. et al. Single-neuronal elements of speech production in humans. Nature (2024).

[49] Dockes, J., et al. NeuroQuery, comprehensive meta-analysis of human brain mapping. eLife 9, e53385 (2020).

[50] Oganian, Y. & Chang, E. F. A speech envelope landmark for syllable encoding in human superior temporal gyrus. Science advances 5, eaay6279 (2019).

[51] Churchland, M. M. et al. Neural population dynamics during reaching. Nature 487, 51–56 (2012).

[52] Crone, N. E., Boatman, D., Gordon, B. & Hao, L. Induced electrocorticographic gamma activity during auditory perception. Clinical Neurophysiology 112, 565–582 (2001).

[53] Lynott, D., Connell, L., Brysbaert, M., Brand, J. & Carney, J. The Lancaster Sensorimotor Norms: Multidimensional measures of Perceptual and Action Strength for 40,000 English words (2019).

[54] Arnal, L. H. & Giraud, A.-L. Cortical oscillations and sensory predictions. Trends in Cognitive Sciences 16, 390–398 (2012).

[55] Fontolan, L., Morillon, B., Liegeois-Chauvel, C. & Giraud, A.-L. The contribution of frequency-specific activity to hierarchical information processing in the human auditory cortex. Nature Communications 5, 4694 (2014).

[56] Leszczyński, M., et al. Dissociation of broadband high-frequency activity and neuronal firing in the neocortex. Science Advances 6, eabb0977 (2020).

[57] Marslen-Wilson, W. D. Functional parallelism in spoken word-recognition. Cognition 25, 71–102 (1987).

[58] Norris, D. & McQueen, J. M. Shortlist B: A Bayesian model of continuous speech recognition. Psychological Review 115, 357–395 (2008).

[59] Lόpez Zunini, R. A., Baart, M., Samuel, A. G. & Armstrong, B. C. Lexical access versus lexical decision processes for auditory, visual, and audiovisual items: Insights from behavioral and neural measures. Neuropsychologia 137, 107305 (2020).

[60] Bentin, S., McCARTHY, GREGORY. & Wood, C. C. Event-related potentials, lexical decision and semantic priming. Electroencephalography and clinical Neurophysiology 60, 1985 (1985).

[61] Barber, H. A., Otten, L. J., Kousta, S.-T. & Vigliocco, G. Concreteness in word processing: ERP and behavioral effects in a lexical decision task. Brain and Language 125, 47–53 (2013).

[62] Vignali, L. et al. Spatiotemporal dynamics of abstract and concrete semantic representations. Brain and Language 243, 105298 (2023).

[63] Rahimi, S., Farahibozorg, S.-R., Jackson, R. & Hauk, O. Task modulation of spatiotemporal dynamics in semantic brain networks: An EEG/MEG study. NeuroImage 246, 118768 (2022).

[64] Lau, E. F., Phillips, C. & Poeppel, D. A cortical network for semantics: (de)constructing the N400. Nature Reviews Neuroscience 9, 920–933 (2008).

[65] Kutas, M. & Federmeier, K. D. Electrophysiology reveals semantic memory use in language comprehension. Trends in Cognitive Sciences 4, 463–470 (2000).

[66] Söderström, P. & Cutler, A. Early neuro-electric indication of lexical match in English spoken-word recognition. PLOS ONE 18, e0285286 (2023).

[67] Hunter, C. R. Early effects of neighborhood density and phonotactic probability of spoken words on event-related potentials. Brain and Language 127, 463–474 (2013).

[68] Winsler, K., Midgley, K. J., Grainger, J. & Holcomb, P. J. An electrophysiological megastudy of spoken word recognition. Language, Cognition and Neuroscience 33, 1063–1082 (2018).

[69] DiCarlo, J. J. & Cox, D. D. Untangling invariant object recognition. Trends in Cognitive Sciences 11, 333–341 (2007).

[70] Rutten, S., Santoro, R., Hervais-Adelman, A., Formisano, E. & Golestani, N. Cortical encoding of speech enhances task-relevant acoustic information. Nature Human Behaviour 3, 974–987 (2019).

[71] Sjerps, M. J., Fox, N. P., Johnson, K. & Chang, E. F. Speaker-normalized sound representations in the human auditory cortex. Nature Communications 10 (2019).

[72] Dykstra, A. R. et al. Individualized localization and cortical surface-based registration of intracranial electrodes. NeuroImage 59, 3563–3570 (2012).

[73] Quiroga, R. Q., Nadasdy, Z. & Ben-Shaul, Y. Unsupervised spike detection and sorting with wavelets and superparamagnetic clustering. Neural computation 16, 1661–1687 (2004).

[74] Boersma, P. Praat, a system for doing phonetics by computer. Glot. Int. 5, 341–345 (2001).

[75] Kisler, T., Reichel, U. & Schiel, F. Multilingual processing of speech via web services. Computer Speech & Language 45, 326–347 (2017).

[76] Crosse, M. J., Di Liberto, G. M., Bednar, A. & Lalor, E. C. The multivariate temporal response function (mTRF) toolbox: A MATLAB toolbox for relating neural signals to continuous stimuli. Frontiers in human neuroscience 10, 604 (2016).

[77] Maris, E. & Oostenveld, R. Nonparametric statistical testing of EEG- and MEG-data. Journal of Neuroscience Methods 164, 177–190 (2007).

[78] Pesaran, B. et al. Investigating large-scale brain dynamics using field potential recordings: Analysis and interpretation. Nature Neuroscience (2018).

[79] Barnett, L. & Seth, A. K. The MVGC multivariate Granger causality toolbox: A new approach to Granger-causal inference. Journal of Neuroscience Methods 223, 50–68 (2014).

[80] Barnett, L. & Seth, A. K. Granger causality for state-space models. Physical Review E 91, 040101 (2015).

