## Supplementary Material for "Top-down semantic predictions align phonetic neuronal dynamics in human superior temporal gyrus"

### Phonetic and semantic mTRF features

**Supplementary Table 1 | Vowels used in the mTRF models for the semantic categorization task.** Duration is computed as the mean in milliseconds across all occurrences of the same phoneme.

| | *First formant* | | *Second formant* | | | *Task: participants 1 and 2* | | *Speech: participant 1* | | *Speech: participant 3* | |
|---|---|---|---|---|---|---|---|---|---|---|---|
| **Vowel** | **Low** | **High** | **Front** | **Middle** | **Back** | **Repetitions** | **Duration** | **Repetitions** | **Duration** | **Repetitions** | **Duration** |
| /ae/ | 1 | 0 | 1 | 0 | 0 | 64 | 87.8 | 34 | 142.9 | 90 | 117.4 |
| /ʌ/ | 1 | 0 | 0 | 0 | 1 | 103 | 47.0 | 165 | 60.7 | 246 | 55.0 |
| /ɪ/ | 0 | 1 | 1 | 0 | 0 | 61 | 56.1 | 116 | 77.6 | 205 | 84.1 |
| /ɜ/ | 1 | 0 | 0 | 1 | 0 | 43 | 89.4 | 43 | 93.0 | 61 | 66.0 |
| /uː/ | 0 | 1 | 0 | 0 | 1 | 31 | 81.8 | 30 | 114.0 | 50 | 129.5 |
| /ɒ/ | 1 | 0 | 0 | 0 | 1 | 49 | 73.4 | 53 | 104.2 | 68 | 72.4 |
| /ɛ/ | 1 | 0 | 1 | 0 | 0 | 44 | 66.4 | 63 | 94.6 | 121 | 118.0 |
| /ɔ/ | 1 | 0 | 0 | 0 | 1 | 29 | 91.5 | 29 | 138.9 | 44 | 112.7 |
| /i/ | 0 | 1 | 1 | 0 | 0 | 55 | 89.6 | 77 | 112.8 | 150 | 108.4 |
| /u/ | 0 | 1 | 0 | 0 | 1 | 7 | 57.1 | 13 | 78.1 | 14 | 64.3 |
| /ʊ/ | 0 | 1 | 0 | 0 | 1 | 8 | 33.8 | 7 | 102.9 | 23 | 78.3 |

**Supplementary Table 2** | **Consonants used in the mTRF models.** Duration is computed as the mean in milliseconds across all occurrences of the same phoneme.

| | *Articulation place* | | | | | | *Articulation manner* | | | | | *Task: participants 1 and 2* | | *Speech: participant 1* | | *Speech: participant 3* | |
|---|---|---|---|---|---|---|---|---|---|---|---|---|---|---|---|---|---|
| **Consonant** | **Bilabial** | **Labiodental** | **Alveolar** | **Velar** | **Uvular** | **Glottal** | **Plosive** | **Nasal** | **Fricative** | **Approximant** | **Lateral approximant** | **Repetitions** | **Duration** | **Repetitions** | **Duration** | **Repetitions** | **Duration** |
| /ŋ/ | 0 | 0 | 0 | 1 | 0 | 0 | 0 | 1 | 0 | 0 | 0 | 19 | 52.8 | 26 | 65.4 | 51 | 82.1 |
| /k/ | 0 | 0 | 0 | 1 | 0 | 0 | 1 | 0 | 0 | 0 | 0 | 135 | 41.0 | 43 | 41.4 | 105 | 67.0 |
| /l/ | 0 | 0 | 1 | 0 | 0 | 0 | 0 | 0 | 0 | 0 | 1 | 123 | 47.9 | 76 | 73.9 | 117 | 58.4 |
| /t/ | 0 | 0 | 1 | 0 | 0 | 0 | 1 | 0 | 0 | 0 | 0 | 120 | 53.0 | 83 | 46.0 | 223 | 101.9 |
| /n/ | 0 | 0 | 1 | 0 | 0 | 0 | 0 | 1 | 0 | 0 | 0 | 89 | 46.2 | 123 | 118.0 | 223 | 125.1 |
| /p/ | 1 | 0 | 0 | 0 | 0 | 0 | 1 | 0 | 0 | 0 | 0 | 75 | 36.7 | 22 | 43.2 | 61 | 67.7 |
| /f/ | 0 | 1 | 0 | 0 | 0 | 0 | 0 | 0 | 1 | 0 | 0 | 45 | 36.7 | 26 | 33.2 | 42 | 120.8 |
| /d/ | 0 | 0 | 1 | 0 | 0 | 0 | 1 | 0 | 0 | 0 | 0 | 51 | 34.0 | 39 | 40.5 | 109 | 66.9 |
| /ʁ/ | 0 | 0 | 0 | 0 | 1 | 0 | 0 | 0 | 1 | 0 | 0 | 100 | 38.3 | 64 | 53.8 | 96 | 34.8 |
| /ʃ/ | 0 | 0 | 1 | 0 | 0 | 0 | 0 | 0 | 1 | 0 | 0 | 22 | 70.7 | 25 | 106.5 | 24 | 110.4 |
| /s/ | 0 | 0 | 1 | 0 | 0 | 0 | 0 | 0 | 1 | 0 | 0 | 104 | 65.0 | 108 | 112.8 | 151 | 102.0 |
| /b/ | 1 | 0 | 0 | 0 | 0 | 0 | 1 | 0 | 0 | 0 | 0 | 79 | 22.0 | 13 | 39.2 | 64 | 40.2 |
| /dʑ/ | 0 | 0 | 1 | 0 | 0 | 0 | 0 | 0 | 1 | 0 | 0 | 18 | 59.0 | 10 | 63.0 | 20 | 82.0 |
| /m/ | 1 | 0 | 0 | 0 | 0 | 0 | 0 | 1 | 0 | 0 | 0 | 60 | 46.6 | 46 | 64.9 | 104 | 181.2 |
| /g/ | 0 | 0 | 0 | 1 | 0 | 0 | 1 | 0 | 0 | 0 | 0 | 36 | 35.0 | 34 | 60.3 | 52 | 40.8 |
| /v/ | 0 | 1 | 0 | 0 | 0 | 0 | 0 | 0 | 1 | 0 | 0 | 13 | 28.5 | 24 | 45.5 | 38 | 84.9 |
| /h/ | 0 | 0 | 0 | 0 | 0 | 1 | 0 | 0 | 1 | 0 | 0 | 25 | 37.9 | 27 | 41.2 | 39 | 49.3 |
| /z/ | 0 | 0 | 1 | 0 | 0 | 0 | 0 | 0 | 1 | 0 | 0 | 13 | 50.0 | 55 | 111.2 | 91 | 129.8 |
| /tɕ/ | 0 | 0 | 1 | 0 | 0 | 0 | 0 | 0 | 1 | 0 | 0 | 31 | 67.8 | 70 | 41.4 | 136 | 54.5 |
| /w/ | 0 | 0 | 0 | 1 | 0 | 0 | 0 | 0 | 0 | 1 | 0 | 22 | 35.8 | 70 | 58.5 | 97 | 41.5 |
| /θ/ | 0 | 0 | 1 | 0 | 0 | 0 | 0 | 0 | 1 | 0 | 0 | 10 | 29.0 | 19 | 34.4 | 39 | 159.4 |

**Supplementary Table 3 | Semantic features for the semantic categorization task (participants 1 and 2).**

| | Semantic features | Repetitions |
|---|---|---|
| *Conceptual category* | Object | 200 |
| | Animal | 200 |
| *Perceptual category* | Bigger than a foot | 200 |
| | Smaller than a foot | 200 |
| *Semantic decision (participant 1)* | Bigger than a foot | 139 |
| | Smaller than a foot | 261 |
| *Semantic decision (participant 2)* | Bigger than a foot | 224 |
| | Smaller than a foot | 176 |

**Supplementary Table 4 | Lexical semantics feature for natural speech perception - participants 1 and 3.** Values represent average strength for each Lancaster norm for each participant across all words.

| Lancaster sensorimotor norm | Participant 1 (664 words) | Participant 3 (1137 words) |
|---|---|---|
| Auditory | 1.83 | 1.85 |
| Gustatory | 0.36 | 0.41 |
| Haptic | 1.13 | 1.17 |
| Interoceptive | 1.2 | 1.31 |
| Olfactory | 0.44 | 0.54 |
| Visual | 2.73 | 2.77 |
| Foot, leg | 1.03 | 1.04 |
| Hand, arm | 1.49 | 1.52 |
| Head | 2.37 | 2.50 |
| Mouth | 1.4 | 1.52 |
| Torso | 0.91 | 0.95 |

### Raster plots, mTRF models for phonetic groups, and PCA details

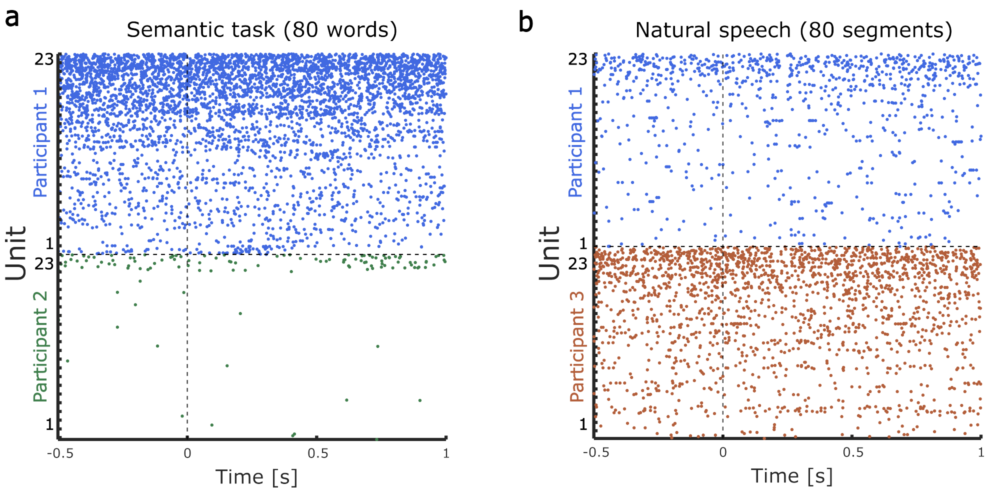


**Supplementary Fig. 1** | **Raster plots a.** Raster plots for the semantic-task participants (1 and 2) for 23 most-spiking units and for 80 random words. Time 0 indicates word onset and all words were 0.5 seconds long. Colors indicate participants as in the main Fig. 1. Despite sparse spiking in participant 2, we still observed the same population effects as for participant 1 (main Fig. 2). **b.** Raster plots for the natural-speech participants (1 and 3) for 23-most spiking units and for 80 random speech segments of the same duration as in the semantic task (1.5 seconds). Time 0 indicates word onset during natural speech, but each word was of a different length, and there was speech before and after the word during the 1.5-second period. Colors indicate participants as in the main Fig. 3.

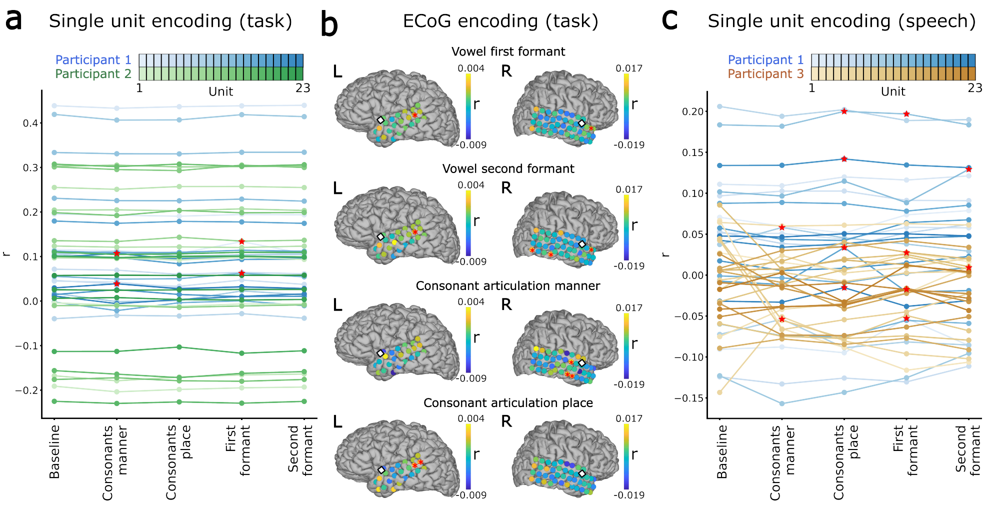


**Supplementary Fig. 2 | Phonetic encoding at the single-unit and LFP level in the aSTG.** **a.** Pearson correlation coefficient (r values) for models including each phonetic feature group fitted to each of the 23 most-spiking single units in the ensemble for the semantic-task participants 1 (blue) and 2 (green). For both participants, the lines for different units are color-coded based on firing rate, from lower (light) to higher (dark). Red stars indicate units for which r values of the fitted models are significantly higher ($p < 0.05$) than the chance level. **b** Encoding of vowel first formant, vowel second formant, consonant manner of articulation, and consonant place of articulation across ECoG channels in the two semantic-task participants. Colors indicate differences in r values compared to the baseline model. Red stars indicate significance as above. Squares indicate MEAs. **c.** Pearson correlation coefficient (r values) for models including each phonetic feature group fitted to each of the 23 most-spiking single units in the ensemble for the natural speech participants 1 (blue) and 3 (orange). For both participants, the lines for different units are color-coded based on firing rate, from lower (light) to higher (dark). Red stars indicate units for which r values of the fitted models are significantly higher ($p < 0.05$) than the chance level.

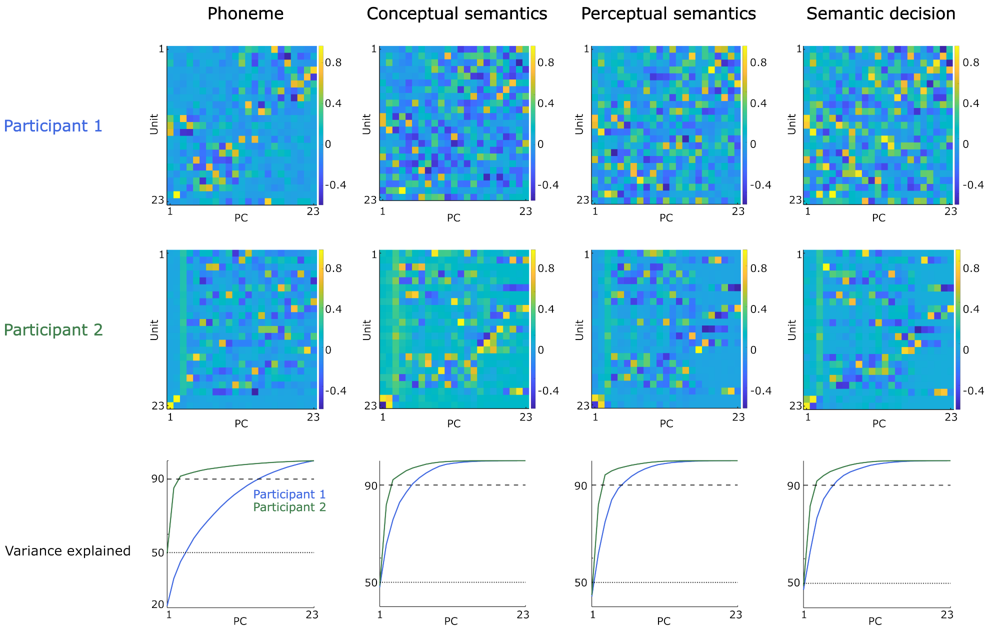


**Supplementary Fig. 3** | **Semantic categorization task PCA** Principal component (PC) coefficients of isolated single units sorted by increasing firing rate for Participant 1 (first row) and Participant 2 (second row), and variance explained (third row) for phonetic and semantic kernels used in the semantic task. Each column indicates the PCs for different mTRF feature kernels from main Fig. 2a, respectively phoneme, conceptual semantics, perceptual semantics, and semantic decision. For Participant 1, several units are represented in the first PCs, indicating distributed encoding across the 23 units. For Participant 2, despite most variance being explained by the 2 most-spiking units, distributed encoding of all units showed population effects (main Fig 2c and 2d).

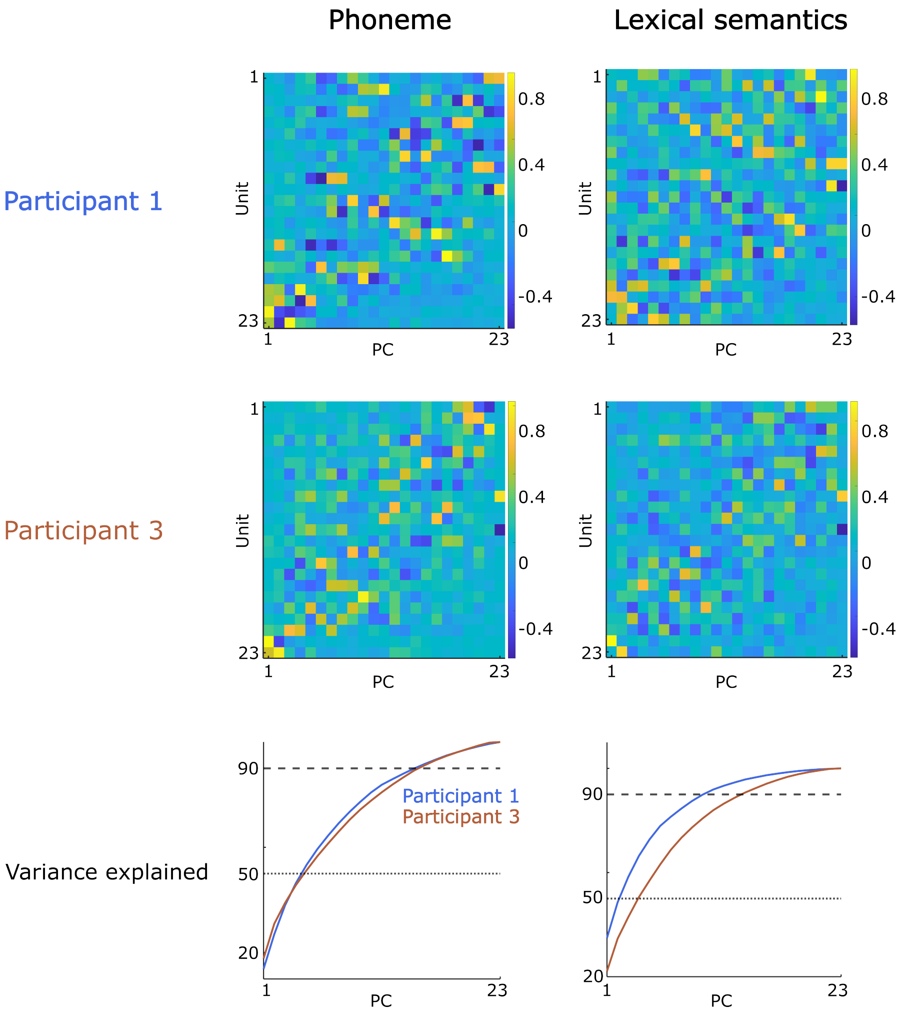


**Supplementary Fig. 4** | **Natural speech PCA** Principal component (PC) coefficients of isolated single units sorted increasingly by firing rate for Participant 1 (first row) and Participant 3 (second row), and variance explained (third row) for phonetic and semantic kernels used in natural speech. Each column indicates the PCs for different mTRF feature kernels, respectively phoneme (main Fig. 3b) and lexical semantics (main Fig. 3d). Distributed encoding is apparent in both participants for both features, and reflected in population effects (main Fig 3c and 3d).

### Clustering control analyses for the semantic task

To confirm our clustering results, we conducted several control analyses. We first confirmed that our results did not change when using PC spaces with more than two PC dimensions. Thus, we computed the clustering index for all four phonetic features (vowel first formant, vowel second formant, consonant manner of articulation, consonant place of articulation - Supplementary Figs. 5 and 6) and semantic features (perceptual, conceptual, decision - Supplementary Figs. 7 and 8) up to the sixth PC dimension. The results observed for two dimensions largely generalized when using more dimensions for the PC space.

For the vowel second formant group, we additionally ascertained that the lack of clustering was not caused by the fact that it contained three phonetic features (front, middle, and back position of the tongue) instead of two as for the first formant group (high, low). We replicated the lack of clustering effects by using two instead of three phonetic features (front and back position of the tongue, Supplementary Fig. 9).

To confirm the observed clustering of phonemes, we performed three alternative analyses: (i) linear discriminant analysis (LDA) classifier (Supplementary Fig. 10); (ii) for vowels, rank regression analysis (rank regression was performed to test whether vowels not only separate, but are also ordered by their first formant value in the PC space, Supplementary Fig. 11); (iii) k-means clustering (Supplementary Fig. 12). To reduce the number of plots, these alternative analyses are shown only for Participant 1, but the same was observed for Participant 2.

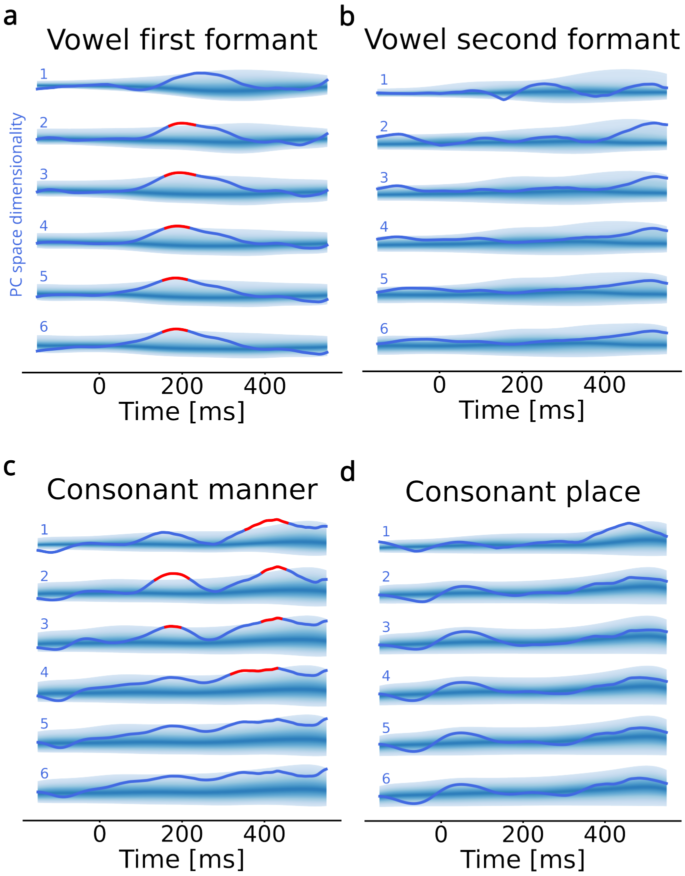


**Supplementary Fig. 5 | Clustering index for phonetic features across dimensions during semantic categorization task in Participant 1. a.** Clustering index for vowels grouped by first formant (high and low tongue position) up to six PC dimensions. The 95% confidence region of the surrogate (chance-level) distribution is shown with brighter shading for increasingly peripheral percentiles. Red segments indicate significant periods after multiple comparison correction (cluster-based test). Numbers indicate the dimensionality of the PC space. **b.** Clustering index for vowels grouped by second formant (front to back tongue position) up to six PC dimensions. Coloring and numbering as in a. **c.** Clustering index for consonants grouped by manner of articulation (plosive, nasal, fricative, approximant, and lateral approximant) up to six PC dimensions. Coloring and numbering as in a. **d.** Clustering index for consonants grouped by place of articulation (bilabial, labiodental, dental, alveolar, velar, uvular, and glottal) up to six PC dimensions. Coloring and numbering as in a.

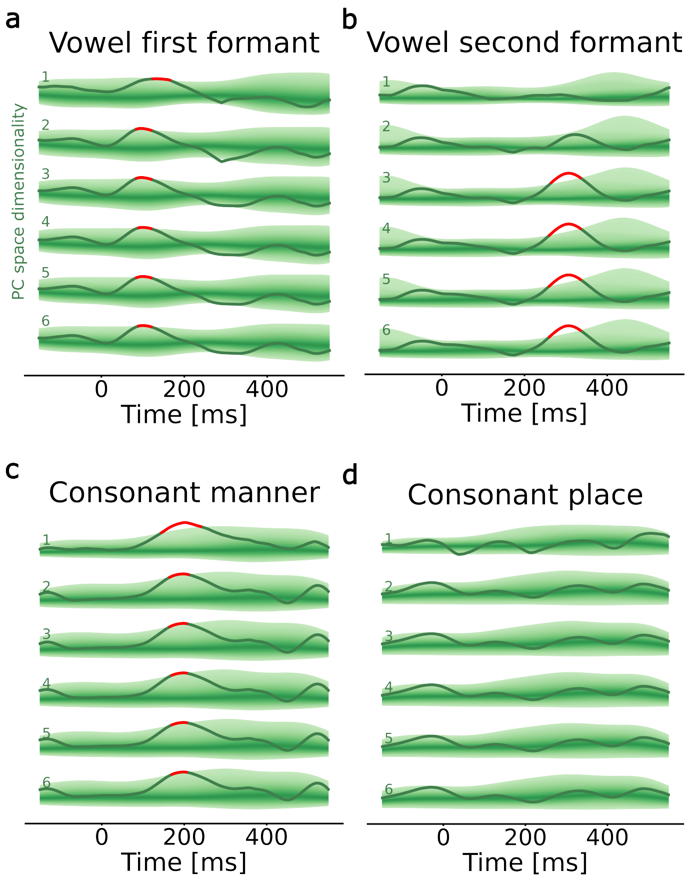


**Supplementary Fig. 6 | Clustering index for phonetic features across dimensions during semantic categorization task in Participant 2. a.** Clustering index for vowels grouped by first formant (high and low tongue position) up to six PC dimensions. The 95% confidence region of the surrogate (chance-level) distribution is shown with brighter shading for increasingly peripheral percentiles. Red segments indicate significant periods after multiple comparison correction (cluster-based test). Numbers indicate the dimensionality of the PC space. **b.** Clustering index for vowels grouped by second formant (front to back tongue position) up to six PC dimensions. Coloring and numbering as in a. **c.** Clustering index for consonants grouped by manner of articulation (plosive, nasal, fricative, approximant, and lateral approximant) up to six PC dimensions. Coloring and numbering as in a. **d.** Clustering index for consonants grouped by place of articulation (bilabial, labiodental, dental, alveolar, velar, uvular, and glottal) up to six PC dimensions. Coloring and numbering as in a.

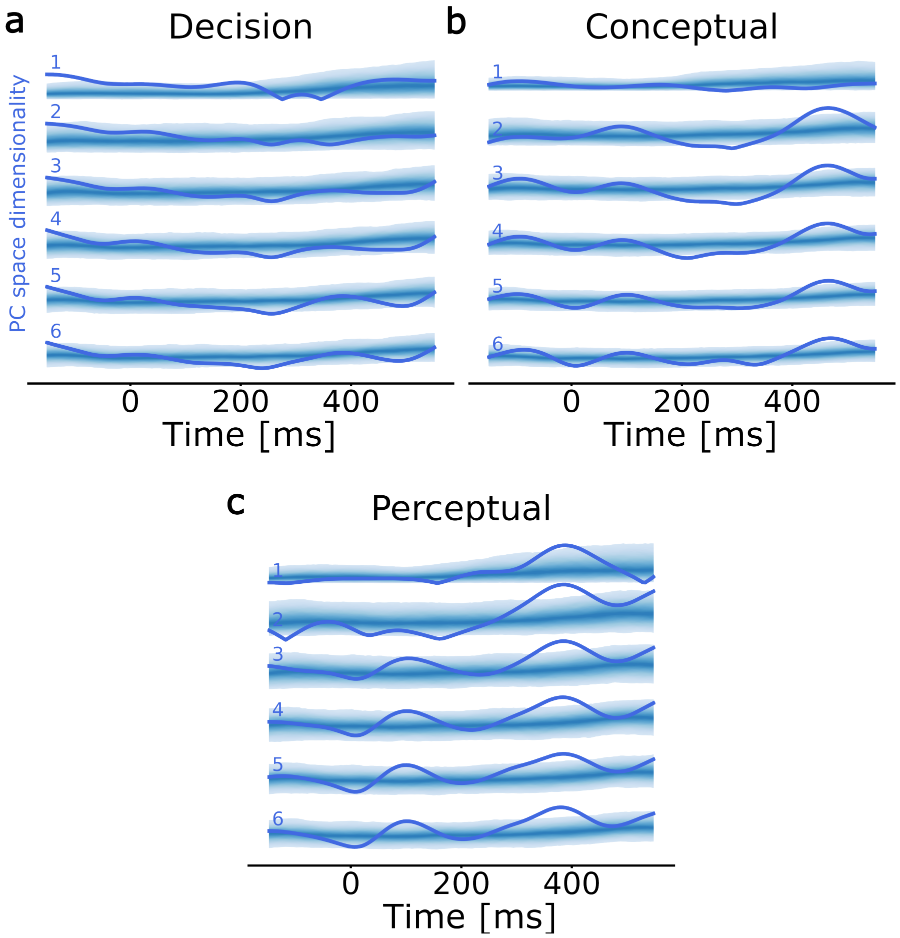


**Supplementary Fig. 7 | Clustering index for semantic features across dimensions during semantic categorization task in Participant 1. a.** Clustering index for semantic decision (response bigger vs. smaller) kernels. The 95% confidence region of the surrogate (chance-level) distribution is shown with brighter shading for increasingly peripheral percentiles. **b.** Clustering index for conceptual semantics (object vs. animal). Coloring and numbering as in a. **c.** Clustering index for perceptual semantics (bigger vs. smaller). Coloring and numbering as in a.

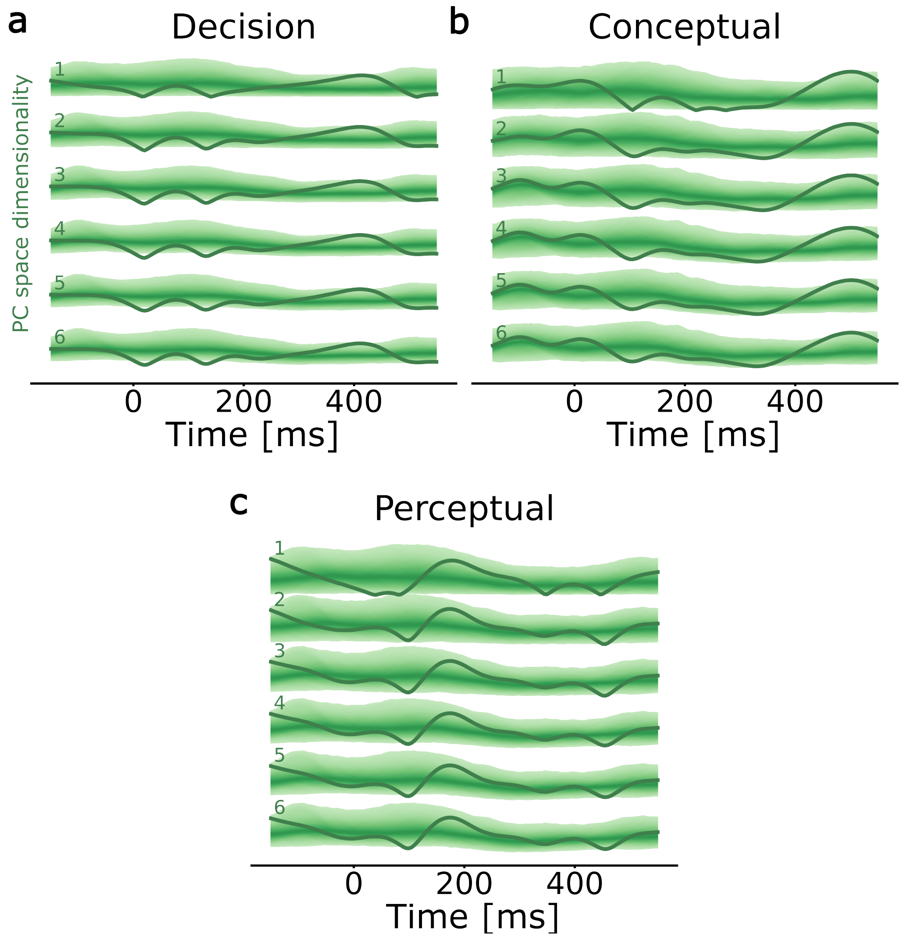


**Supplementary Fig. 8 | Clustering index for semantic features across dimensions during semantic categorization task in Participant 2. a.** Clustering index for semantic decision (response bigger vs. smaller) kernels. The 95% confidence region of the surrogate (chance-level) distribution is shown with brighter shading for increasingly peripheral percentiles. **b.** Clustering index for conceptual semantics (object vs. animal). Coloring and numbering as in a. **c.** Clustering index for perceptual semantics (bigger vs. smaller). Coloring and numbering as in a.

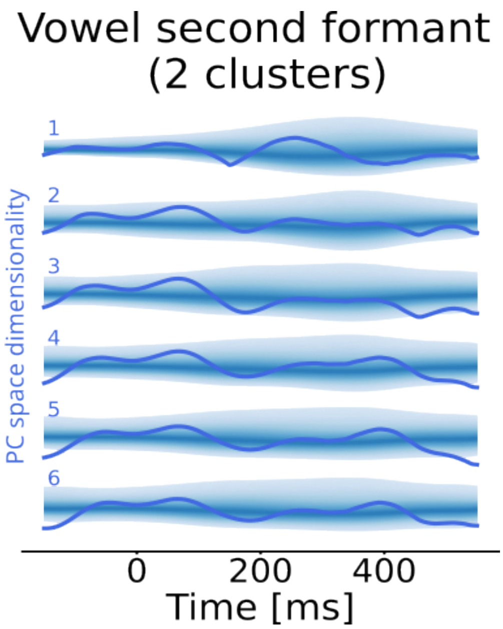


**Supplementary Fig. 9 | Clustering for the vowel second formant group using only two phonetic features in Participant 1.** The 95% confidence region of the surrogate (chance-level) distribution is shown with brighter shading for increasingly peripheral percentiles.

#### Linear discriminant analysis (LDA) classifier

We ran the LDA classifier for each of the 4 phonetic groups (vowel first formant, vowel second formant, consonant manner, consonant place). For each time point, we first ran an LDA classifier which is a Gaussian mixture model to compute the means of the multivariate normal distributions for each class. Then, we computed the average Euclidean distance between all class means and compared it against the distribution of 1000 surrogates. This is similar to the between-cluster distances from our clustering algorithm, with the difference that here the class mean (cluster centroid) is a parameter of the estimated multivariate distribution, and not computed directly by averaging class elements.

Replicating the clustering index analyses, at 200 ms we observed a significant separation of vowels into first formant categories (Supplementary Fig. 10a). Linear separators estimated at 0 and 200 ms are shown in Supplementary Fig. 10b. No peaks was observed for the vowel second formant (Supplementary Fig. 10c). LDA also indicated the two previously observed peaks for consonant manner (200 and 400 ms), however, here, they did not reach significance (Supplementary Fig. 10d). Finally, LDA revealed a significant peak for consonant place at 50 ms (Supplementary Fig. 10e) that was not observed before.

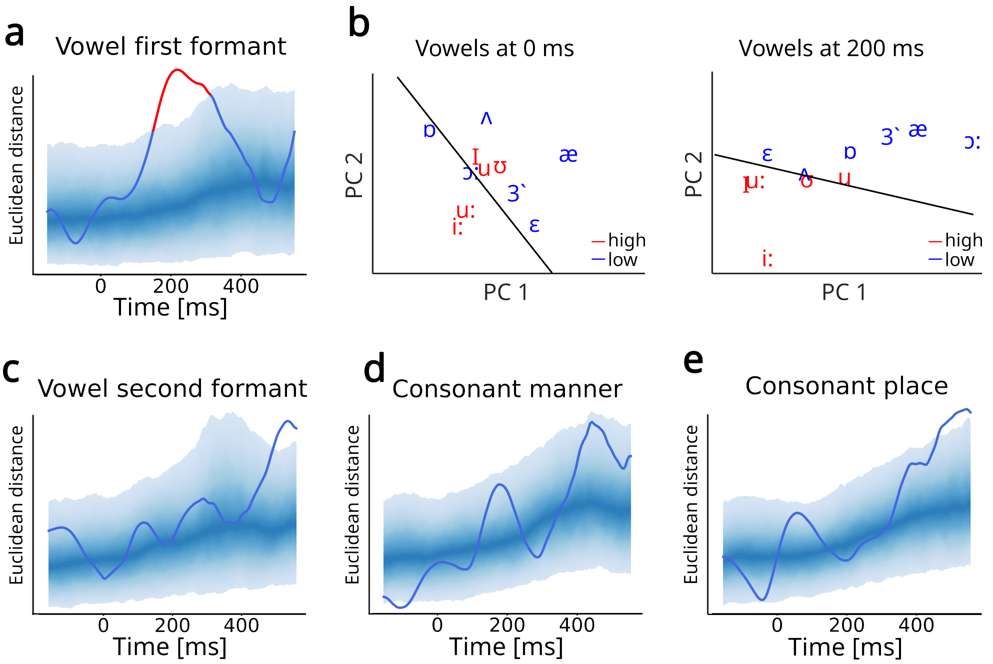


**Supplementary Fig. 10 | Linear discriminant analysis during the semantic categorization task in Participant 1. a.** Vowel first formant. The 95% confidence region of the surrogate (chance-level) distribution is shown with brighter shading for increasingly peripheral percentiles. Red segments indicate significant periods after multiple comparison correction (cluster-based test). **b.** Estimated linear separators for vowel first formant feature at 0 and 200 ms. **c** Vowel second formant. Shading and red segments are as in a. **d.** Consonant manner of articulation. Coloring as in a. **e.** Consonant place of articulation. Coloring as in a.

#### Rank regression

Since formants are actually ordered by their frequency values along the first and second formant axis, we additionally explored whether the actual ordering of formant frequency values was encoded in the low-dimensional space. To that aim, we ran a rank regression analysis, where a rank value (1-7) was assigned to each vowel, based on the standard IPA table. At each time point, the ranked order of vowels was correlated with their coordinates on the first three axes (PC1, PC2, and PC3), and compared against a distribution of 1000 surrogates.

We observed a significant ordering of vowels based on their first formant ranks on all three PCs (Fig. 11a). We performed the same regression for the second formant values and observed a significant correlation at about 0 ms, but only on PC1 (Supplementary Fig. 11b). As the two consonant groups are based on categorical and not continuous variables, rank regression analysis is not applicable.

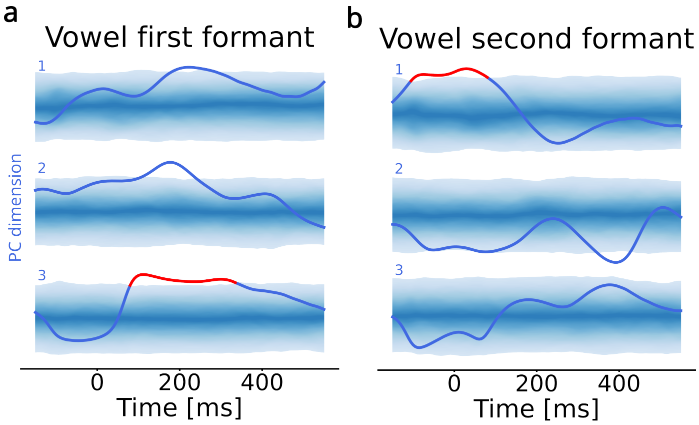


**Supplementary Fig. 11 | Rank regression during semantic categorization in Participant 1. a.** Correlation between vowel first formant ranks and coordinates on PC axes. The 95% confidence region of the surrogate (chance-level) distribution is shown with brighter shading for increasingly peripheral percentiles. Red segments indicate significant periods after multiple comparison correction (cluster-based test). Numbers indicate PC dimensions. **b.** Correlation between vowel second formant ranks and coordinates on PC axes. Coloring and numbering as in **a**.

### K-means clustering

We used k-means clustering to investigate whether the same clustering of phonemes would emerge in a data-driven fashion. Our clustering results of the main text necessitate assigning a priori each of the N phonemes to a cluster (e.g., for the vowel first formant feature see Supplementary Fig. 12a), and the clustering index is then quantified based on the Euclidean distances of those a priori chosen clusters. On the contrary, k-means clustering is an unsupervised method that partitions all N phonemes into k clusters based on their proximity in PC space. Thus, for each time point, we first ran k-means clustering 1000 times, as the clustering results might change based on the algorithm's random initialization. Then, we computed an average N-by-N attribution matrix that indicated how often each of the N phonemes was clustered together (Supplementary Fig. 12b). Finally, we correlated the resulting k-means attribution matrix with the attribution matrix of the actual, linguistically-based clusters, and compared the correlation value against the distribution of correlation values for the 1000 surrogates at each time point (Supplementary Fig. 12c-f).

Replicating previous clustering results for the vowel first formant group, this analysis revealed a significant period at about 200 ms during which data-driven clusters overlapped with the linguistic ones (Supplementary Fig. 12c). This analysis further revealed a significant peak at 75 ms for vowel second formant that was not present before (Supplementary Fig. 12d). The 200-ms peak was also observed again for the consonant manner group, with an additional peak at about 0 ms, and no significant peak at 400 ms (Supplementary Fig. 12e). No significant peaks were observed for consonant place (Supplementary Fig. 12f).

Despite its advantageous data-driven perspective, there are also some important drawbacks of this method, that can be observed in the shape of the correlation curve for the vowel first formant group (Supplementary Fig. 12c). Namely, at 200 ms there is a

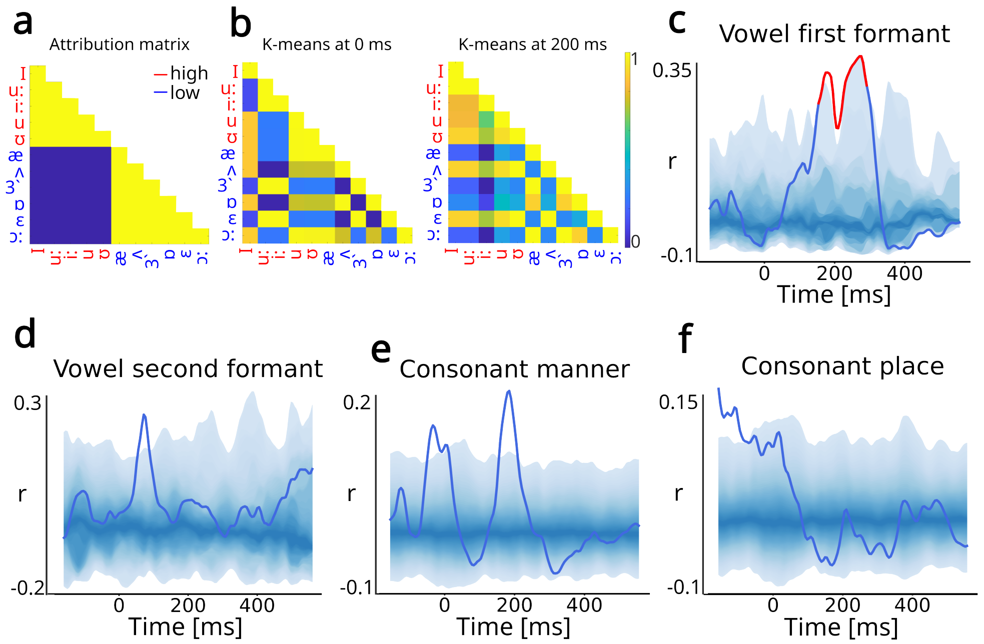


**Supplementary Fig. 12 | K-means clustering during semantic categorization in Participant 1. a.** Attribution matrix for the vowel first formant feature. **b.** K-means matrices for the vowel first formant feature at 0 and 200 ms. **c.** Correlation between vowel first formant attribution matrix (**a**) and corresponding k-means matrices (**b**) at each time point. The 95% confidence region of the surrogate (chance-level) distribution is shown with brighter shading for increasingly peripheral percentiles. Red segments indicate significant periods after multiple comparison correction (cluster-based test). **d.** Correlation between vowel second formant and corresponding k-means attribution matrices at each time point. Coloring as in **c**. **e.** Correlation between consonant manner and corresponding k-means matrices at each time point. Coloring as in **c**. **f.** Correlation between consonant place and corresponding k-means matrices at each time point. Coloring as in **c**.

decrease in the correlation value (although still significant), because at that time point vowels /ʌ/ and /ɛ/, which belong to the 'low' group, are closer in space to the 'high' group (compare the 200-ms insets on Fig. 2e and Supplementary Fig. 12b). However, it is apparent that, despite their spatial proximity to the 'high' group, vowels /ʌ/ and /ɛ/ can be attributed to the 'low' group because the two clusters can be linearly separated right next to the positions of these two vowels in space (Supplementary Fig. 10b).

#### Summary of control analyses

Supplementary Table 5 summarizes all peaks observed across different analyses for all 4 groups. Although some analyses sporadically revealed significant peaks for vowel second formant (k-means) and consonant place (LDA) groups, the peaks that were consistent across all analyses are the same peaks as observed in our initial clustering analysis: 200 ms for vowel first formant and consonant manner, and 400 ms for consonant manner alone.

**Supplementary Table 5 | Summary of the significant window locations during semantic categorization identified through control analyses.** Bold entries indicate peaks that were consistent across analyses. Entries in brackets indicate existing peaks that did not cross the significance threshold. Entries in red indicate peaks surviving multiple comparison correction (cluster-based test). All values are in ms.

| **Phonetic feature group** | **Clustering** | **LDA** | **Rank regression** | **K-means** |
|---|---|---|---|---|
| Vowel first formant | **200** | **200** | **200** | **200** |
| Vowel second formant | | | 0 | 75 |
| Consonant manner | **200, 400** | **(200, 400)** | not applicable | 0, **200** |
| Consonant place | | 450 | not applicable | |

### Clustering control analyses for natural speech perception

We performed the same control analyses for natural speech perception: generalization of clustering for up to 6-dimensional PC spaces for phonetic (Supplementary Figs. 13 and 14) and semantic features (Supplementary Fig. 15). As for the semantic task, for phonetic features we also performed LDA (Supplementary Fig. 16), rank regression (Supplementary Fig. 17), and correlation with K-means attribution (Supplementary Fig. 18). We then summarized the significant peaks observed during natural speech perception across all analyses. (Supplementary Table 6).

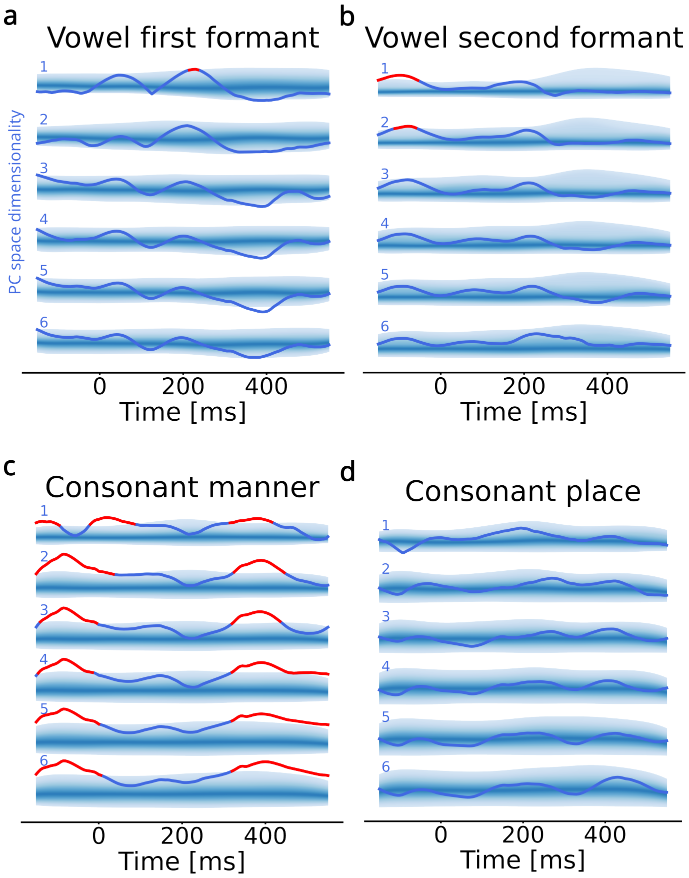


**Supplementary Fig. 13 | Clustering index across dimensions during natural speech perception in Participant 1. a.** Clustering index for vowels grouped by the first formant (high and low tongue position) up to 6 PC dimensions. The 95% confidence region of the surrogate (chance-level) distribution is shown with brighter shading for increasingly peripheral percentiles. Numbers indicate the dimensionality of the PC space. **b.** Clustering index for vowels grouped by the second formant up to six PC dimensions. Coloring and numbering as in a. **c.** Clustering index for consonants grouped by the manner of articulation up to six PC dimensions. Shading and numbering as in a. **d.** Clustering index for consonants grouped by the place of articulation up to six PC dimensions. Coloring and numbering as in a.

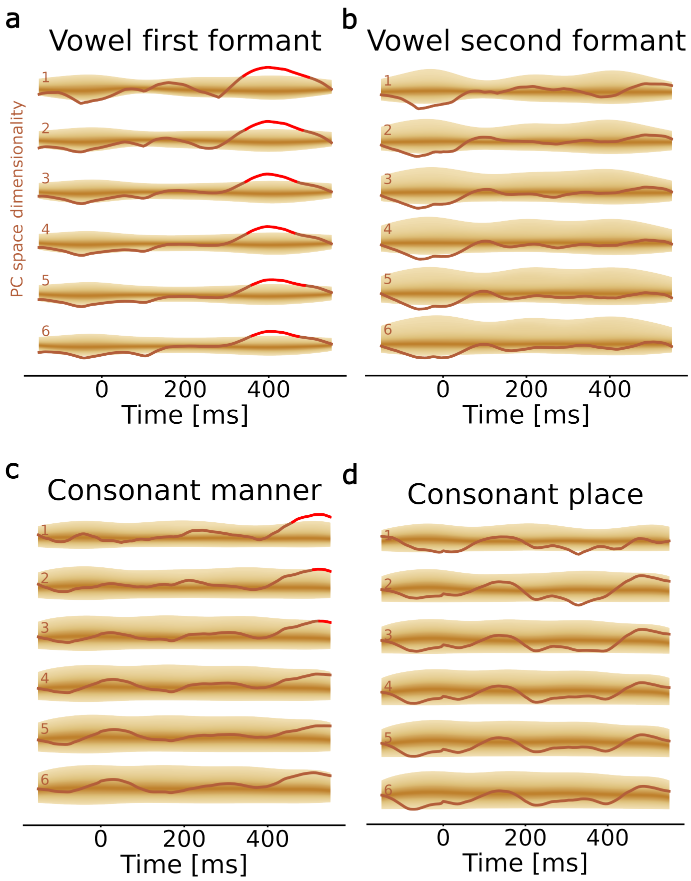


**Supplementary Fig. 14 | Clustering index across dimensions during natural speech perception in Participant 3. a.** Clustering index for vowels grouped by the first formant (high and low tongue position) up to 6 PC dimensions. The 95% confidence region of the surrogate (chance-level) distribution is shown with brighter shading for increasingly peripheral percentiles. Numbers indicate the dimensionality of the PC space. **b.** Clustering index for vowels grouped by the second formant up to six PC dimensions. Coloring and numbering as in a. **c.** Clustering index for consonants grouped by the manner of articulation up to six PC dimensions. Shading and numbering as in a. **d.** Clustering index for consonants grouped by the place of articulation up to six PC dimensions. Coloring and numbering as in a.

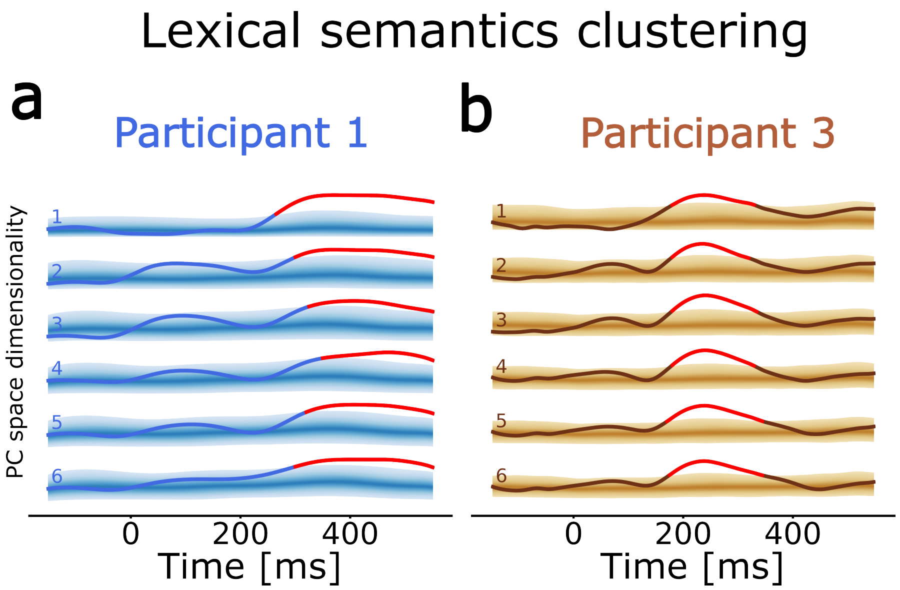


**Supplementary Fig. 15 | Clustering of semantic features across dimensions during natural speech perception. a.** Clustering index for lexical semantics feature in Participant 1. The 95% confidence region of the surrogate (chance-level) distribution is shown with brighter shading for increasingly peripheral percentiles. Red segments indicate significant periods after multiple comparison correction (cluster-based test). Numbers indicate the dimensionality of the PC space. **b.** Clustering index for lexical semantics feature in Participant 3.

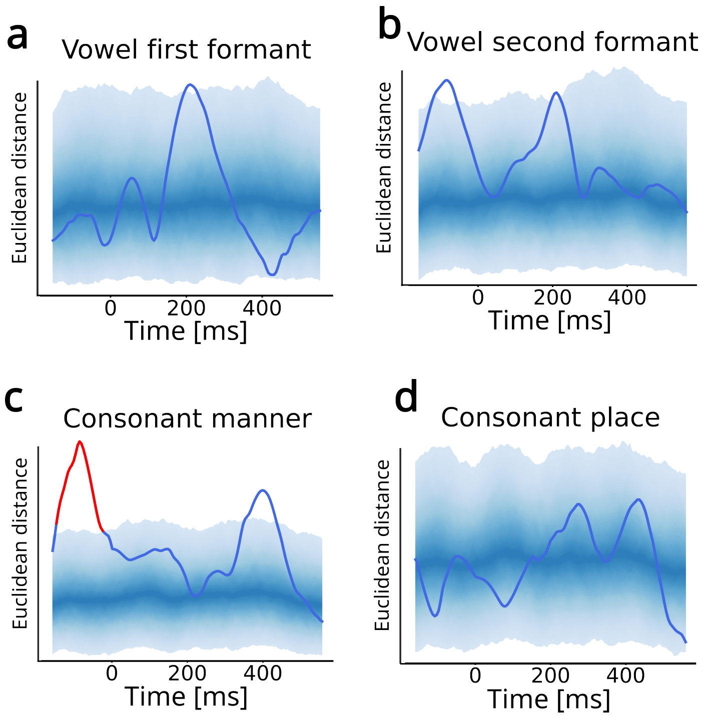


**Supplementary Fig. 16 | Linear discriminant analysis during natural speech perception in Participant 1. a.** Vowel first formant. The 95% confidence region of the surrogate (chance-level) distribution is shown with brighter shading for increasingly peripheral percentiles. Red segments indicate significant periods after multiple comparison correction (cluster-based test). **b.** Vowel second formant. Coloring as in **a**. **c.** Consonant manner of articulation. Coloring as in **a**. **d.** Consonant place of articulation. Coloring as in **a**.

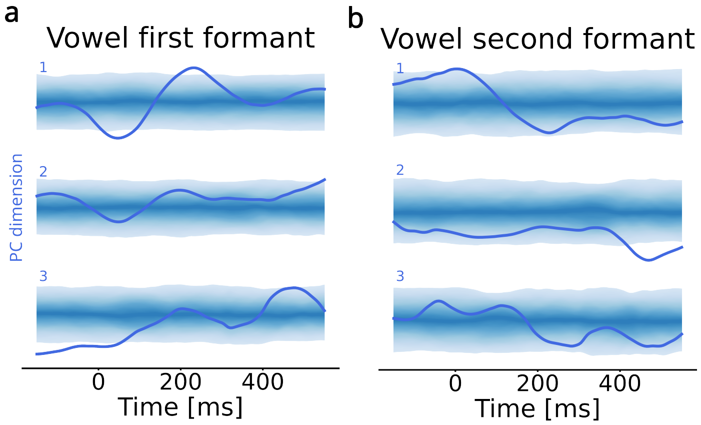


**Supplementary Fig. 17 | Rank regression during natural speech perception in Participant 1. a.** Correlation between vowel first formant ranks and coordinates on PC axes. The 95% confidence region of the surrogate (chance-level) distribution is shown with brighter shading for increasingly peripheral percentiles. Numbers indicate PC dimensions. **b.** Correlation between vowel second formant ranks and coordinates on PC axes. Coloring and numbering as in **a**.

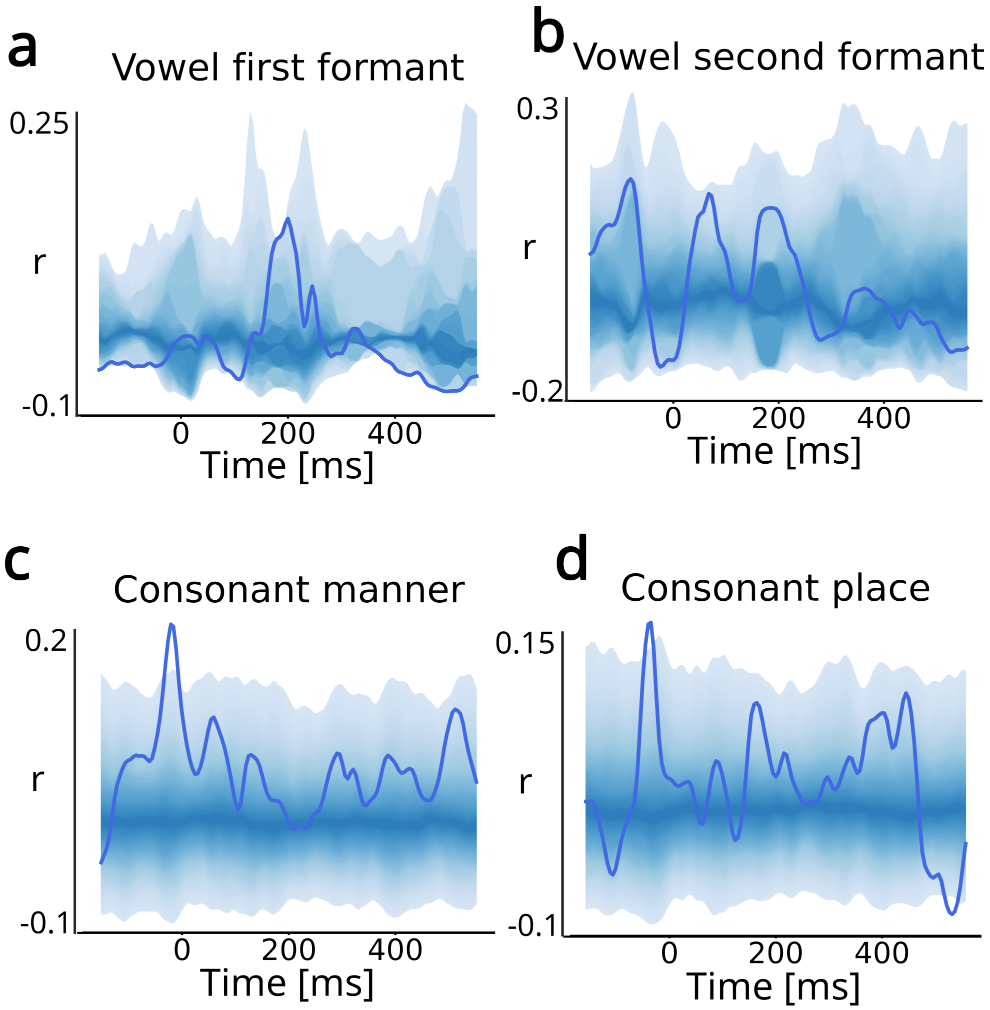


**Supplementary Fig. 18 | K-means clustering during natural speech perception in Participant 1. a.** Correlation between vowel first formant and corresponding k-means attribution matrices at each time point. The 95% confidence region of the surrogate (chance-level) distribution is shown with brighter shading for increasingly peripheral percentiles. **b.** Correlation between vowel second formant and corresponding k-means attribution matrices at each time point. Coloring as in **a**. **c.** Correlation between consonant manner and corresponding k-means attribution matrices at each time point. Coloring as in **a**. **d.** Correlation between consonant place and corresponding k-means attribution matrices at each time point. Coloring as in **a**.

**Supplementary Table 6 | Summary of the significant window locations during natural speech perception identified through control analyses.** Bold entries indicate peaks that were consistent across analyses. Entries in brackets indicate existing peaks that did not cross the significance threshold. Entries in red indicate peaks surviving multiple comparison correction (cluster-based test). All values are in ms.

| **Phonetic feature group** | **Clustering** | **LDA** | **Rank regression** | **K-means** |
|---|---|---|---|---|
| Vowel first formant | (**200**) | **200** | **200** | **200** |
| Vowel second formant | **-100**, (**200**) | **-100, 200** | 0 | |
| Consonant manner | **-100, 400** | **-100, 400** | not applicable | -25 |
| Consonant place | | | not applicable | -25 |

### Top-down semantic and phonetic alignment

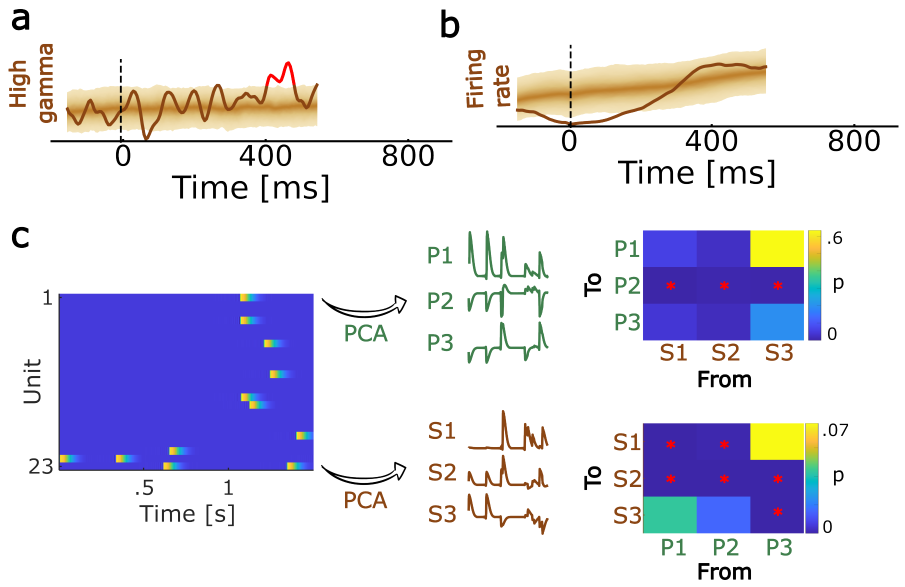


**Supplementary Fig. 19 | Control analyses for the top-down alignment in Participant 1. a.** mTRF kernel for the broadband high-frequency activity (BHA, 70-150 Hz) across the MEA showed a significant peak at around 400 ms, overlapping with the significant periods of the lexical semantic firing rate kernel (main Fig. 4b), and the beta-band word-onset kernel (main Fig. 4c). The 95% confidence region of the surrogate (chance-level) distribution is shown with brighter shading for increasingly peripheral percentiles. Red segments indicate periods surviving multiple comparison correction. **b.** mTRF firing rate kernel aligned to word onset did not have a significant peak at 400 ms, suggesting that the observed BHA peak is not resulting from the underlying neuronal spiking. Coloring as in a. **c.** Granger causality between phonetic and semantic low-dimensional representations. The left plot shows a 1.5-second snippet of the spiking data recorded during natural speech perception. Spiking activity was smoothed with a 25-ms half-Gaussian kernel to obtain a causal firing rate estimates. These are then projected onto three PCs to obtain time-varying phonetic (P) and semantic (S) features, which are then used in the Granger causality analysis (MVGC toolbox). Matrices indicate p-values of Granger's F-test for a causal relationship from the dimensions indicated on the x-axis to the dimensions indicated on the y-axis. The stars indicate significant relationships at the 0.05 threshold. While there were significant causal relationships in both directions (i.e. phonetic to semantic and vice versa), there were more causal relationships in the top-down direction.
